# Degraded sensory coding in a mouse model of *Scn2a-*related disorder and its rescue by CRISPRa gene activation

**DOI:** 10.64898/2025.12.13.694145

**Authors:** Kaeli Vandemark, Hannah R. Monday, Lucia Rodriguez, Manya Zhao, Meeseo Lee, Amit Ahituv, Anwita Satapathy, Nicholas F. Page, Kelly An, Elizabeth Hamada, Nadav Ahituv, Kevin J. Bender, Daniel E. Feldman

## Abstract

Heterozygous loss-of-function mutations in *SCN2A*, a sodium channel gene expressed in cortical pyramidal (PYR) cells, lead to a neurodevelopmental disorder characterized by autism, intellectual disability, and cortical sensory dysfunction. In *Scn2a+/−* mice, PYR dendritic excitability and synapses are impaired, but cortical information processing deficits are unknown. In the whisker somatosensory cortex, we found strongly degraded somatotopic tuning of PYR cells, profoundly blurred whisker maps, and impaired population coding, despite normal overall firing rates. This constitutes a robust biomarker for *Scn2a*-related cortical dysfunction. Parvalbumin (PV) interneurons were also unexpectedly hypofunctional. We tested for rescue of coding deficits in post-critical period, young adult mice by viral delivery of CRISPR activation (CRISPRa) reagents that upregulate *Scn2a*. CRISPRa treatment normalized cortical sensory representations at the single-unit and map levels. This suggests that therapy to increase *Scn2a* expression may be effective in normalizing cortical function in *Scn2a* loss-of-function disorder, even in older children or adults.

## INTRODUCTION

*Scn2a*-related disorders are a family of severe neurodevelopmental disorders (NDDs) caused by mutations in the *Scn2a* gene, which encodes the voltage-gated sodium channel NaV1.2. Heterozygous loss-of-function mutations in *Scn2a* are associated with autism spectrum disorder (ASD), intellectual disability (ID) and developmental delay^1–3^, childhood-onset seizures^4^, and often with cortical visual impairment^5,6^. *Scn2a* is the highest-risk gene identified via exome sequencing in people with ASD^7–10^. Additional sensory features of *Scn2a* loss-of-function mutations include poor sensory registration and muted responses to aversive stimuli^4,11^. These sensory impairments emerge during early childhood^12^, consistent with a neurodevelopmental origin, and may involve dysfunction in sensory cortex, as suggested for other forms of ASD^13–15^. But the nature of sensory cortex dysfunction in *Scn2a*-related disorders is unknown, as is whether, and when, cortical function can be rescued therapeutically.

*Scn2a* critically regulates dendritic excitability in neocortical pyramidal (PYR) cells^16^. In *Scn2a* heterozygous knockout (*Scn2a+/−*) mice, which model *Scn2a* loss-of-function, PYR cells show reduced dendritic excitability, impaired long-term synaptic potentiation, and weak glutamatergic synapses^17–20^ suggesting impaired activity-dependent synaptic maturation. How these cellular-level physiological deficits drive dysfunction at the neural coding and information processing level is unknown. The apparent immaturity of synapses in *Scn2a+/−* mice suggests that neural coding in sensory cortex may be weak or imprecise. Such a coding deficit would align with the sensory behavioral deficits in people, and could provide an effective preclinical biomarker for development of therapeutics to normalize cortical processing. Immature *Scn2a+/−* synaptic phenotypes suggest a failure of early critical period synaptic refinement, and that restoration of dendritic excitability, even in adults, could re-enable normal synaptic maturation to rescue sensory coding and cortical function.

Atypical sensory processing is a common feature of ASD^21,22^, but there is no robust understanding of how neural coding is disrupted in sensory cortex in ASD-related NDDs. Some theories posit circuit hyperexcitability and excess spiking in sensory cortex, yielding overly intense sensory representations^23,24^ or heightened noise^25–27^. But excess spiking is observed in sensory cortex in only in a few mouse models of ASD-related NDDs (e.g., refs ^28,29^), and instead most models show subtle disruption of other coding properties, including modest blurring of single-neuron sensory tuning or maps^30–33^, impaired sensory adaptation^34^, or abnormal oscillations^35,36^. Hypofunction of parvalbumin (PV) inhibitory interneuron circuits is a common feature across many ASD models and is thought to contribute to abnormal sensory coding^37^. Because PV neurons do not express *Scn2a*^19^, PV hypofunction is not predicted in *Scn2a+/−* mice, suggesting a distinct form of coding impairment may occur.

We characterized neural coding in *Scn2a+/−* mice in whisker primary somatosensory cortex (wS1). Neural circuit function, tactile map organization, and critical periods are well understood in wS1^38–40^, making it a powerful system to identify disease-related cortical phenotypes. wS1 processes touch from the whiskers, which are used analogously to human fingertips, and tactile processing is commonly affected in ASD^13,37,41^. We found that *Scn2a+/−* mice have dramatically degraded tactile coding in wS1, both at the single neuron and population (map) levels, which was markedly greater than the subtle blurring observed in other autism mouse models^29,42–45^. This dramatic blurring suggests an immature, unrefined circuit. We used CRISPR activation (CRISPRa) to increase expression of the remaining functional allele to restore gene expression levels^4,46,47^, a therapeutic strategy that is applicable for genetic disorders caused by heterozygous loss-of-function mutations (including *Scn2a*-related ASD/ID and most syndromic forms of ASD). Applying this CRISPRa approach in young adult mice after the critical period effectively rescued neural coding deficits in sensory cortex. This suggests that adult rescue of cortical sensory function is possible in Scn2a-related ASD/ID.

## RESULTS

### Cellular physiology phenotypes in L2/3 PYR cells in *Scn2a+/−* mice

In *Scn2a+/−* mice, pyramidal (PYR) cells in medial prefrontal cortex (mPFC) and visual cortex show reduced slope and amplitude of somatic action potentials, and weak synapses marked by low AMPA:NMDA ratio and reduced AMPA-mediated miniature excitatory postsynaptic currents (mEPSCs)^19^. These phenotypes reflect the role of NaV1.2 in generation of backpropagating action potentials, which are critical for activity-dependent synapse maturation. To test for these cellular phenotypes in wS1, we performed whole-cell recordings from L2/3 PYR cells in acute wS1 slices from P18-22 mice^48^. Action potentials evoked by current injection just above rheobase had significantly lower rising slope in *Scn2a^+/−^* mice relative to WT, apparent in mean spike waveforms and in phase-plane plots (WT: 470 ± 23.0 mV/ms [mean ± SEM], Het: 339 ± 21.3, p=0.0008, unpaired t-test, sample size for this and all measurements are in **Table S1**). Spike amplitude was also reduced (peak Vm in WT: 35.4 ± 1.1 mV, Het: 28.0 ± 1.6, p=0.0016, Mann-Whitney), with no change in spike threshold (**Fig. 1A-C)**, or V_rest_ and R_input_ (not shown). These are the expected direct effects of reduced NaV1.2 density^19^. AMPA-mediated mEPSCs in L2/3 PYR cells had reduced amplitude (WT: −24.1 ± 1.5 pA, Het: −18.8 ± 0.8, p=0.0062, unpaired t-test) but normal mEPSC frequency (WT: 13.1 ± 1.2 Hz, Het: 15.4 ± 1.4, p=0.028, Mann-Whitney), suggesting weaker functional synapses (**Fig. 1D**). Thus, *Scn2a* haploinsufficiency affects spike shape and synaptic properties in L2/3 PYR cells in S1, similar to other cortical areas^19^.

**Figure 1.**
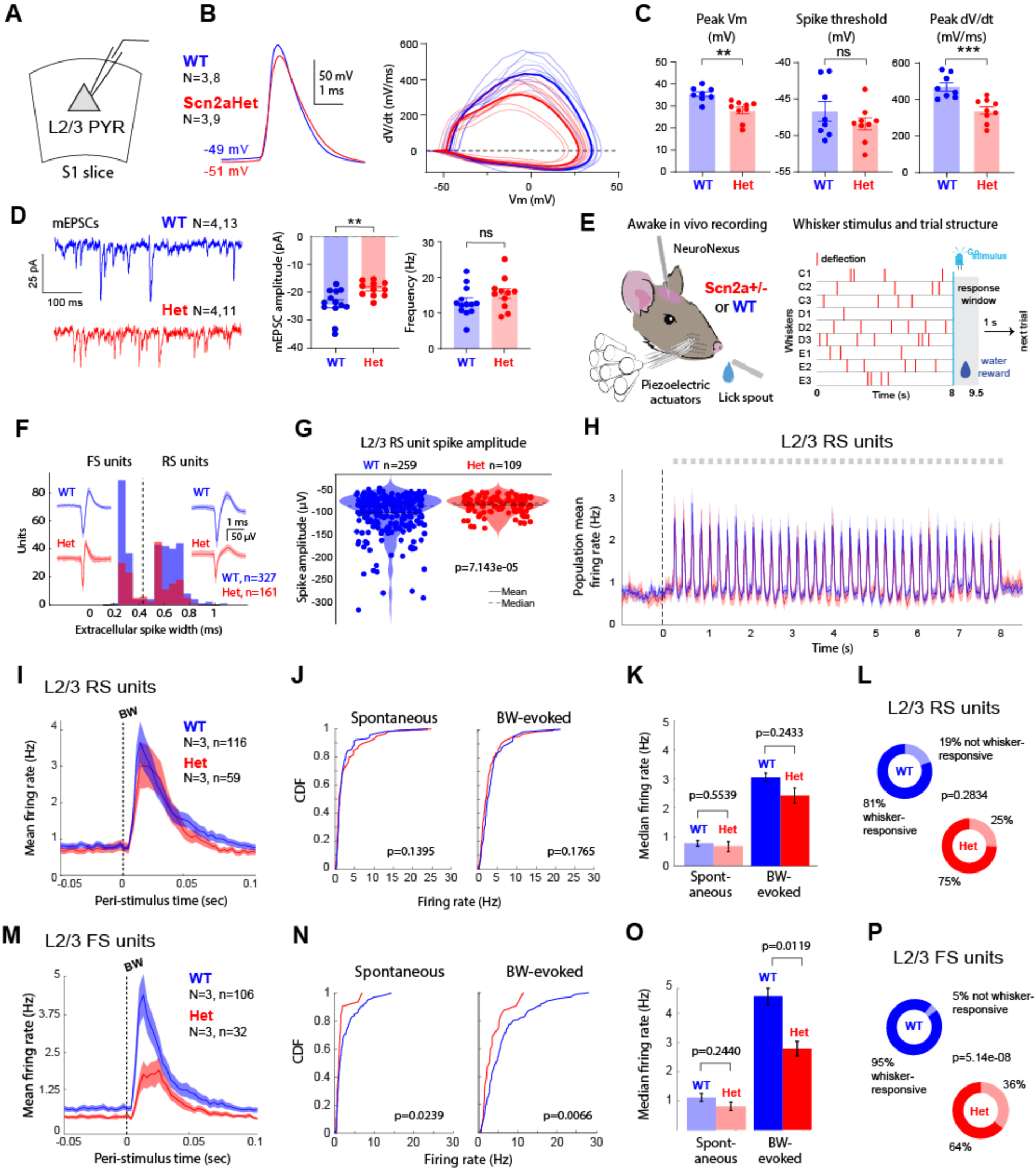
Cellular physiology and in vivo spike rate in L2/3 of Scn2a+/− mice. **A)** Recording from L2/3 pyramidal cells in S1 slices. **B)** Mean action potential waveform (WT: N=3 mice, 8 cells; Het: N=3 mice, 9 cells). Right: Phase-plane plots for the spike of each cell. Thin lines are cells, thick lines are mean. **C**) Peak membrane potential during the spike (Mann-Whitney test, p = 0.0016), spike threshold (unpaired t-test, p = 0.2808), and peak dV/dt for spikes (unpaired t-test, p = 0.0008). **D**) Example mEPSCs from each genotype. Bottom, average mEPSC amplitude per cell (unpaired t-test, p = 0.0062) and mean mEPSC interval per cell (Mann-Whitney test, p = 0.02767). N’s for mEPSC data, WT: N=4 mice, 11 cells, Het: N=4 mice, 13 cells.) **E**) In vivo extracellular recording during behavior, showing trial structure with whisker deflections. **F**) Identification of FS and RS units by spike width. Data are from L2/3 and L4 combined. WT: N=3 mice, n=327 units. *Scn2a*Het: N=3, n=161. Waveforms are mean +- SEM for example units. **G**) Spike amplitude for L2/3 RS units. Each dot is a unit. p-value from Wilcoxon rank-sum test. **H)** Mean population firing rate for L2/3 RS units across all trials. Gray square, whisker deflection (different whiskers interleaved in random order in different trials.). **I-L**) Responses of L2/3 RS units. **I**) Mean population PSTH (WT N= 3 mice, n= 116 units, Het N= 3 mice, n= 59 units). Error shading is bootstrapped 68% CI. **J**) Spontaneous and best whisker (BW)-evoked firing rates. Statistics are from two-sample KS test. **K**) Bootstrapped median firing rates. Error bars are 68% CI. Statistics are from permutation test. **L**) Fraction of units that are whisker-responsive. p-value is from chi-squared goodness-of-fit. **M-P)** Responses of L2/3 FS units. Conventions as in I-L. Alpha was 0.05 for all tests.

### Spontaneous and whisker-evoked spiking in wS1 of awake *Scn2a+/−* mice

To test whether these cellular phenotypes were accompanied by changes in whisker sensory coding in wS1 *in vivo*, we performed dense extracellular spike recordings in wS1 using NeuroNexus probes in awake, head-fixed adult mice (P96-215). Recordings were made in L2/3 and L4 of C2, C1, D2 or B1 whisker columns, verified histologically, as calibrated deflections were applied to 9 whiskers in a 3 x 3 grid centered on the columnar whisker. To establish a uniform behavioral state, mice performed a sensory detection task in which they received water reward for licking in a response window following an occasional LED visual stimulus (flashed in ∼80% of trials). Each trial contained a whisker stimulus period in which single-whisker stimuli (on randomly chosen whiskers) and blanks (no whisker stimulus) were interleaved every 0.2 sec over 8 sec, during which mice were required to withhold licking. This task was chosen because it is simple to train, and allows measurement of whisker responses and tuning properties of single units in wS1 during a task-engaged, lick-free behavioral state (**Fig. 1E**). Task engagement was assessed by comparing lick rates in the response window on LED flash trials (Go trials) vs. no-flash (NoGo) trials. Both genotypes exhibited similar task engagement (**Fig. S1**), and had similar recording locations (**Fig. S2**). After spike sorting, single units were classified as fast-spiking (FS; parvalbumin [PV] interneurons) or regular-spiking (RS; mostly PYR cells) using a spike width criterion that separated two clear peaks in the spike width distribution (**Fig. 1F**). We previously confirmed through opto-tagging that L2/3 FS units identified with this method are PV interneurons^48^.

In *Scn2a+/−* mice, L2/3 RS units had extracellular spike waveforms with lower amplitude than in WT mice (−101.46 µV WT, −88.20 µV *Scn2a+/−*; p=7.143e-5, Wilcoxon rank-sum test) (**Fig. 1G**). This effect was reliable in every mouse (n=3 per genotype), despite the unit-to-unit variability in spike amplitude that reflects distance from the electrode, cell morphology, and other factors (**Fig. S2**). Reduced extracellular spike amplitude is an expected correlate of slower intracellular spike slope (dV/dt), and thus is a biomarker of reduced NaV1.2 density in PYR cells^19^.

At the neural coding level, we first tested the hypothesis that overall spontaneous or sensory-evoked firing rate was altered in wS1. Reduced spiking may be expected in *Scn2a+/−* mice because of reduced mEPSC amplitude (**Fig. 1D**) and more silent synapses^19^. However, when we analyzed all whisker-responsive L2/3 RS units, there was no apparent difference between genotypes in overall mean firing rate across the whisker stimulus period, or in mean sensory adaptation (**Fig. 1H**). Quantitative analysis showed no difference between WT and *Scn2a+/−* mice in spontaneous firing (from blank stimuli: two- sample Kolmogorov-Smirnov [KS] test, p=0.1395; bootstrapped median firing rate [Hz]: 0.76 ± 0.11 WT, 0.83 ± 0.28 *Scn2a+/−*; p=0.5539, permutation test) or in spiking responses to each cell’s best whisker (BW) (KS test, p=0.4974; bootstrapped median firing rate [Hz]: 3.18 ± 0.18 WT, 2.46 ± 0.47 *Scn2a+/−*; p=0.2433) (**Fig. 1I-K**). There was also no difference in the fraction of L2/3 RS units that were whisker-responsive (81% WT, 75% *Scn2a+/−*; p=0.2834, chi-squared test, **Fig. 1L**). Thus, L2/3 RS unit firing rates were not reduced in *Scn2a+/−*mice, nor were they increased as in the cortical hyperexcitability theory of ASD^23,24^.

Surprisingly, L2/3 FS units recorded simultaneously at the same sites had substantially reduced firing in *Scn2a+/−* mice, both for spontaneous activity (KS test, p=0.0239, bootstrapped median firing rate [Hz]: 1.17 ± 0.08 WT, 0.81 ± 0.11 *Scn2a*+/−; p=0.2440, permutation test) and BW-evoked responses (KS test, p=0.0066, bootstrapped median firing rate [Hz]: 4.67 ± 0.32 WT, 2.79 ± 0.29 *Scn2a*+/−; p=0.0119, permutation test) (**Fig. 1M-O**). The fraction of whisker-responsive FS units was also substantially reduced (95% WT, 64% *Scn2a^+/−^*; p=5.1e-8, chi-squared test, **Fig. 1P**). L2/3 FS units are PV interneurons^48^. While PV inhibition is hypofunctional in sensory cortex in many genetic mouse models of ASD^49^, this is unexpected in *Scn2a+/−*, because PV interneurons are not thought to express *Scn2a* (Ref^47^ and see “PV hypofunction”, below). This suggests that reduced PV spiking is an indirect, secondary consequence of *Scn2a* loss in excitatory neurons.

### *Scn2a+/−* mice exhibit profoundly degraded somatotopic tuning in L2/3

Modest blurring of sensory feature tuning or maps is observed in some mouse models of ASD-related NDDs^37,50–52^, and is a sign of degraded information representation in cortex. To test whether *Scn2a^+/−^* mice have blurred sensory tuning, we analyzed single-unit whisker somatotopic tuning across the nine whiskers of the 3x3 piezo array, which were centered on the columnar whisker (CW) for the recorded column in wS1. In the wS1 whisker map, PYR cells in each column are narrowly tuned to the CW or nearby whiskers on the face^38–40^. We analyzed units that were tuned to the CW, because all eight immediate surround whiskers (SWs) were sampled.

In WT mice, L2/3 RS units were narrowly tuned to one or a few whiskers, as normal for S1^39,53–55^. In contrast, in *Scn2a^+/^* mice, most L2/3 RS units were very broadly tuned (**Fig. 2A**). We calculated mean somatotopic tuning curve across L2/3 RS units (**Fig. 2B**). We also constructed a rank-ordered whisker tuning curve for each unit, which assesses tuning width independent from somatotopic structure (**Fig. 2C**). Both analyses revealed markedly broader tuning in *Scn2a^+/−^* mice. Mean rank-ordered tuning curves were sharp for WT units, and significantly broader in *Scn2a^+/−^* mice, with the extent of broadening particularly apparent when tuning curves were normalized to the strongest whisker (**Fig. 2D**). 57% of L2/3 RS units in WT mice had one whisker (the BW) that activated the unit significantly more strongly than any other whisker, while 59% of *Scn2a^+/−^* units had 7-9 statistically equivalent best whiskers (eBWs) (**Fig. 2E**). To quantify tuning width on an individual unit basis, we calculated lifetime sparseness^56^ (whisker selectivity) for each unit. Lifetime sparseness was markedly lower in *Scn2a^+/−^* mice (mean: 0.26 WT, 0.08 *Scn2a^+/−^*, median: 0.20 WT, 0.06 *Scn2a^+/−^*; p=0.0073, permutation test), indicating broader whisker tuning (**Fig. 2F**).

**Figure 2:**
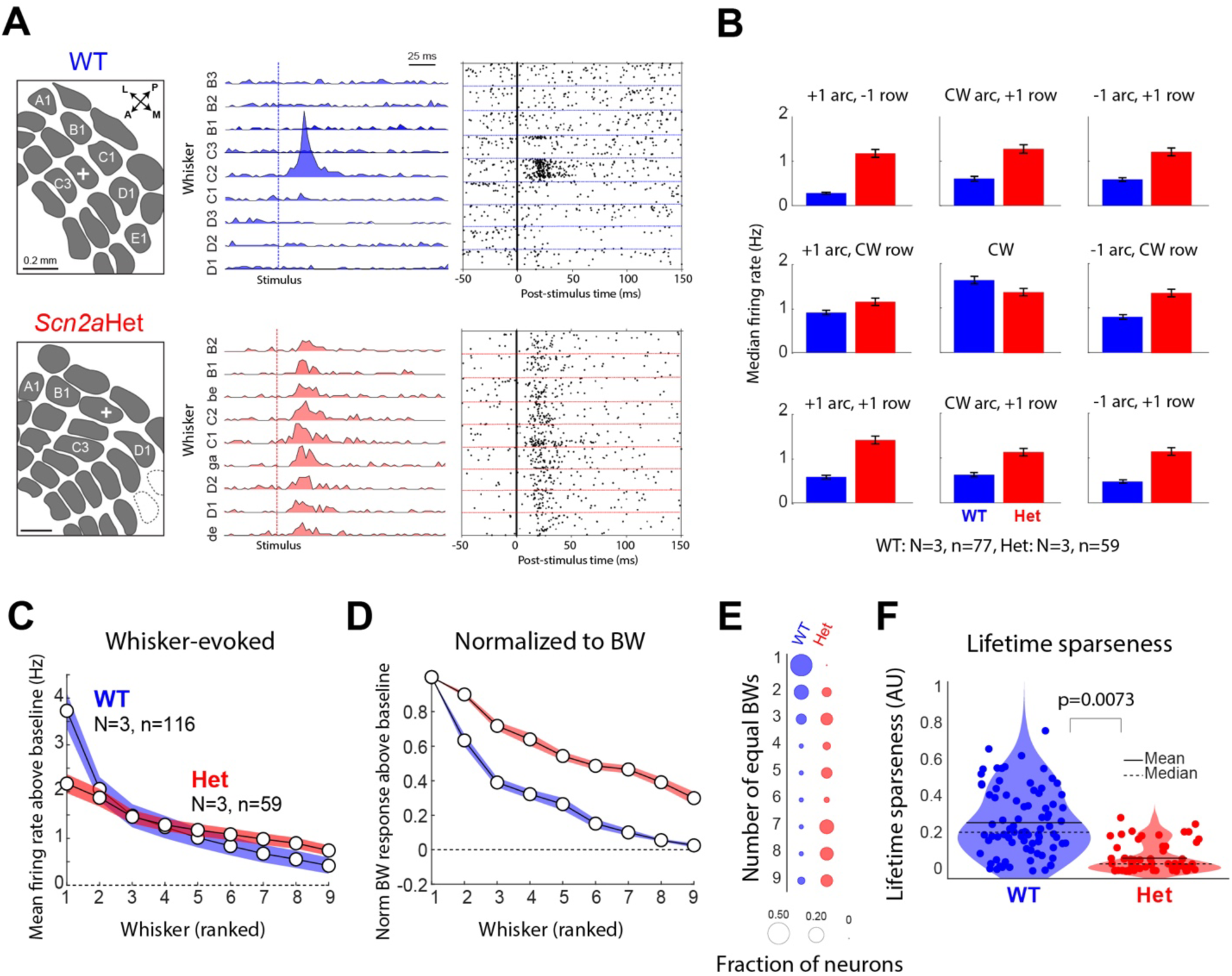
Degraded somatotopic tuning of L2/3 RS units in awake *Scn2a^+/−^* mice. **A)** Representative 9 whisker somatotopic receptive fields for a wild-type and *Scn2a*Het L2/3 RS unit. Left, L2/3 recording site location (cross) in wS1, relative to barrel boundaries in L4 from cytochrome oxidase stain. Dashed outlines, missing barrels. The raster and PSTH are from the same unit. Vertical line: whisker deflection. **B**) Population average somatotopic tuning for L2/3 RS units, for the 3x3 grid of whiskers centered around the CW for the recorded column. Mean bootstrapped firing rate and 68% CI are shown. WT: N=3, n=77 units. Scn2aHet: N=3, n=59 units. **C**) Mean rank-ordered, baseline-subtracted whisker tuning curves for L2/3 RS units. WT: N=3, n=116 units. Scn2aHet: N=3, n=59 units. Dashed line: spontaneous baseline activity. Error shading is 68% CI. **D**) Tuning curves from (C) normalized to each unit’s BW response before averaging. **E**) Number of equivalent BWs across the population. Circle size indicates fraction of units. **F**) Quantification of tuning sharpness by lifetime sparseness. Each dot is one cell. Solid line: mean, dashed line: median. P-value from permutation test. Alpha was 0.05 for all tests.

Broader tuning was observed in L2/3 RS units in every individual *Scn2a+/−* mouse (**Fig. S2C**), and in additional mice recorded with Neuropixels probes (n=1 WT, 1 *Scn2a^+/−^*, not included in the data above) (**Fig. S3**). Broad tuning was not due to differences in recording site location, which was similar between genotypes (**Fig. S2A**). It was also not an inherent feature of low-spike-amplitude L2/3 RS units, because when we sub-selected WT units with low spike amplitudes matched to *Scn2a+/−* RS units, these WT units had the same sharp somatotopic tuning as the full L2/3 RS wild type dataset (**Fig. S2D**). Despite broad tuning, the average spike rate elicited by any whisker stimulus was not significantly increased in *Scn2a+/−*, indicating a redistribution of spikes rather than excess total spiking among L2/3 RS units (**Fig. S2E**).

### Degraded tactile maps

Degraded tuning at the single-unit level in wS1 predicts blurring of population-level somatotopic maps. To test this, we used intrinsic signal optical imaging (ISOI), a hemodynamic measure of neural activity^57–59^. ISOI signals report whisker-evoked functional activation in wS1, originate largely in L2/3^58^, and are centered in the activated whisker’s column^57,60^. In anesthetized mice, we delivered calibrated single-whisker deflection trains on C1, C2, or C3 whiskers, while measuring whisker-evoked intrinsic signal relative to baseline (dR/R, **Fig. 3A**). Activation of each pixel (corresponding to negative dR/R) was normalized to the most strongly activated pixel, and the area of activation was quantified within contours of 0.8, 0.6, or 0.4 of this maximum dR/R value. In a representative WT mouse, C1 whisker stimulation activated a spatially focused area in wS1, with the 0.8 ISOI contour tightly coupled to the C1 whisker anatomical column from post-hoc cytochrome oxidase staining (**Fig. 3B**). In a *Scn2a^+/−^* mouse, C1 deflection activated a pronouncedly broader region that spanned a large set of whisker-related columns (**Fig. 3B**). Across four mice per genotype, C1, C2, and C3 whiskers activated substantially larger areas in wS1 in *Scn2a^+/−^*mice compared to WT (**Fig. 3C**). This broadening of the ISOI area occurred without significant changes in signal amplitude of the most strongly activated pixel (**Fig. S4**). Therefore, functional somatotopic maps are strongly degraded in *Scn2a^+/−^* mice.

**Figure 3:**
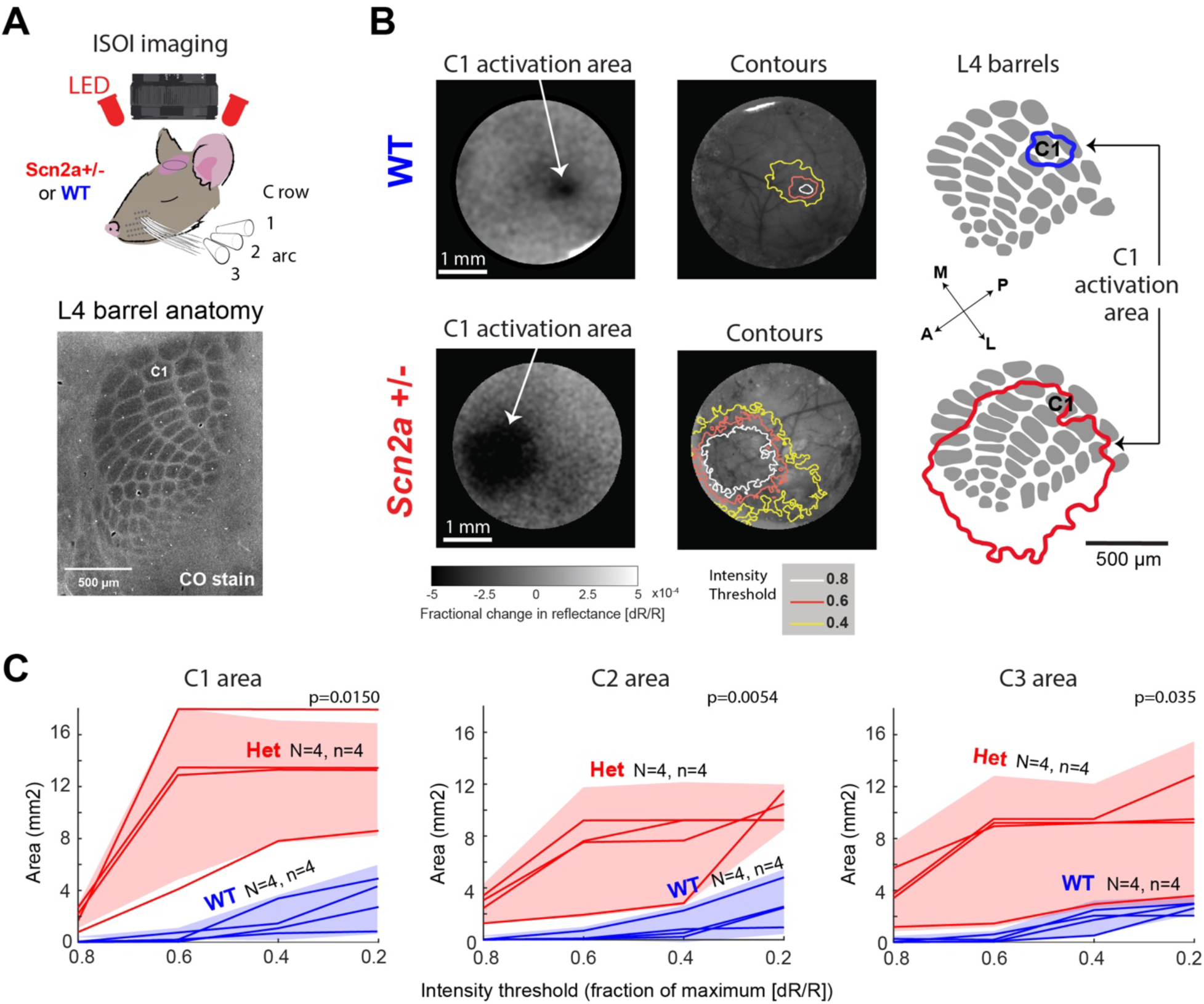
Degraded map somatotopy in *Scn2a^+/−^* mice measured with intrinsic signal optical imaging (ISOI). **A**) Top: Schematic showing measurement of whisker-evoked hemodynamic activity in anesthetized mice using ISOI. A train of single-whisker deflections was delivered to a single whisker (C1, C2 or C3) on each trial. Bottom: cytochrome oxidase (CO)-stained tissue section showing barrel pattern in wS1 of a *Scn2a*Het mouse. **B**) Left: Whisker activation areas for 2 representative mice, showing dR/R evoked by the C1 whisker relative to pre-stimulus baseline, and dR/R contours at 0.8, 0.6, and 0.4 of the maximal dR/R superimposed on the surface vasculature for that mouse. Right: C1 activation area (80% contour, for the same mice as in panel B, left overlaid on barrel pattern. The C1 anatomical column is labeled. **C**) Quantification of whisker activation area for whiskers C1, C2, and C3 in 4 WT mice and 4 Scn2aHet mice. Each line is a mouse; shaded region is 95% CI.

Somatotopic blurring was also evident in the local field potential (LFP). Negative-going LFP transients largely reflect summed synaptic potentials and spiking of local neuronal populations^61–63^. We compared LFP in L2/3 across genotypes in awake, task-engaged mice, simultaneously with the extracellular spike measurements (**Fig. S5A**). From each genotype, 3 mice were recorded with NeuroNexus probes (same recordings as spike data in **Figs. 1-2**), and an additional mouse was recorded with NeuroPixels (same data as **Fig. S3**). We calculated whisker receptive fields from LFP in L2/3, and quantified rank-ordered tuning curves from negative-going LFP amplitude (**Fig. S5C**). Somatotopic tuning measured in the center of the receptive field (whisker ranks 1-4) was broader in *Scn2a^+/−^* mice than in WT, as quantified by a center tuning sharpness index (median: WT 1.47 ± 0.24, Scn2a^+/−^ 0.69 ± 0.31, p=0.0319 (**Fig. S5D**). In contrast, LFP responses evoked by weak, low-rank whiskers (rank 5-8) were not reliably different across the genotypes, so that tuning sharpness across all 9 whiskers, quantified by lifetime sparseness, was not significantly different between genotypes (median: WT 0.52 ± 0.17, *Scn2a^+/−^* 0.30 ± 0.13; p=0.1422, permutation test) (**Fig. S5E**). CW-evoked LFP magnitude did not differ between genotypes, consistent with single-unit results (median: WT −67 ± −10 µV, *Scn2a^+/−^* −55 ± −7; p=0.3082, permutation test) (**Fig. S5F**). Thus, somatotopic tuning of local L2/3 neuronal populations, as assessed by LFP, also showed evidence of degraded tuning, most prominently within tuning curve centers.

### Degraded somatotopic tuning, but intact anatomical tactile maps, in L4

The somatotopic map in wS1 originates in L4, where neurons are clustered into whisker-related barrels and show precise somatotopic tuning^38–40^. To assess whether neural coding was also disrupted in L4, we analyzed L4 RS units recorded in the same penetrations as the L2/3 data above. L4 RS units in *Scn2a+/−* mice showed normal spontaneous and BW-evoked firing rates (spontaneous: KS test, p=0.6407, bootstrapped median firing rate [Hz]: 0.77 ± 0.23 WT, 0.84 ± 0.37 *Scn2a^+/−^*; p=0.6393, permutation test; BW-evoked: KS test, p=0.3673, 3.66 ± 0.28 WT, 3.05 ± 0.40 *Scn2a^+/−^*; p=0.4571, permutation test) (**Fig. 4A-D**), and a normal fraction of whisker-responsive units (84% WT, 73% *Scn2a^+/−^*; p=0.1891, chi-squared test). However, like in L2/3, whisker tuning was markedly broadened for L4 RS units in *Scn2a^+/−^* mice, assessed by mean somatotopic tuning, rank-ordered tuning, and number of equal best whiskers (**Fig. 5E-H**), as well as lifetime sparseness (mean: 0.29 WT, 0.04 *Scn2a^+/−^*, median: 0.22 WT, 0.04 *Scn2a^+/−^*; p=0.0017, permutation test) (**Fig. 5I**). Thus, abnormally broad tuning was also strongly evident in L4.

**Figure 4.**
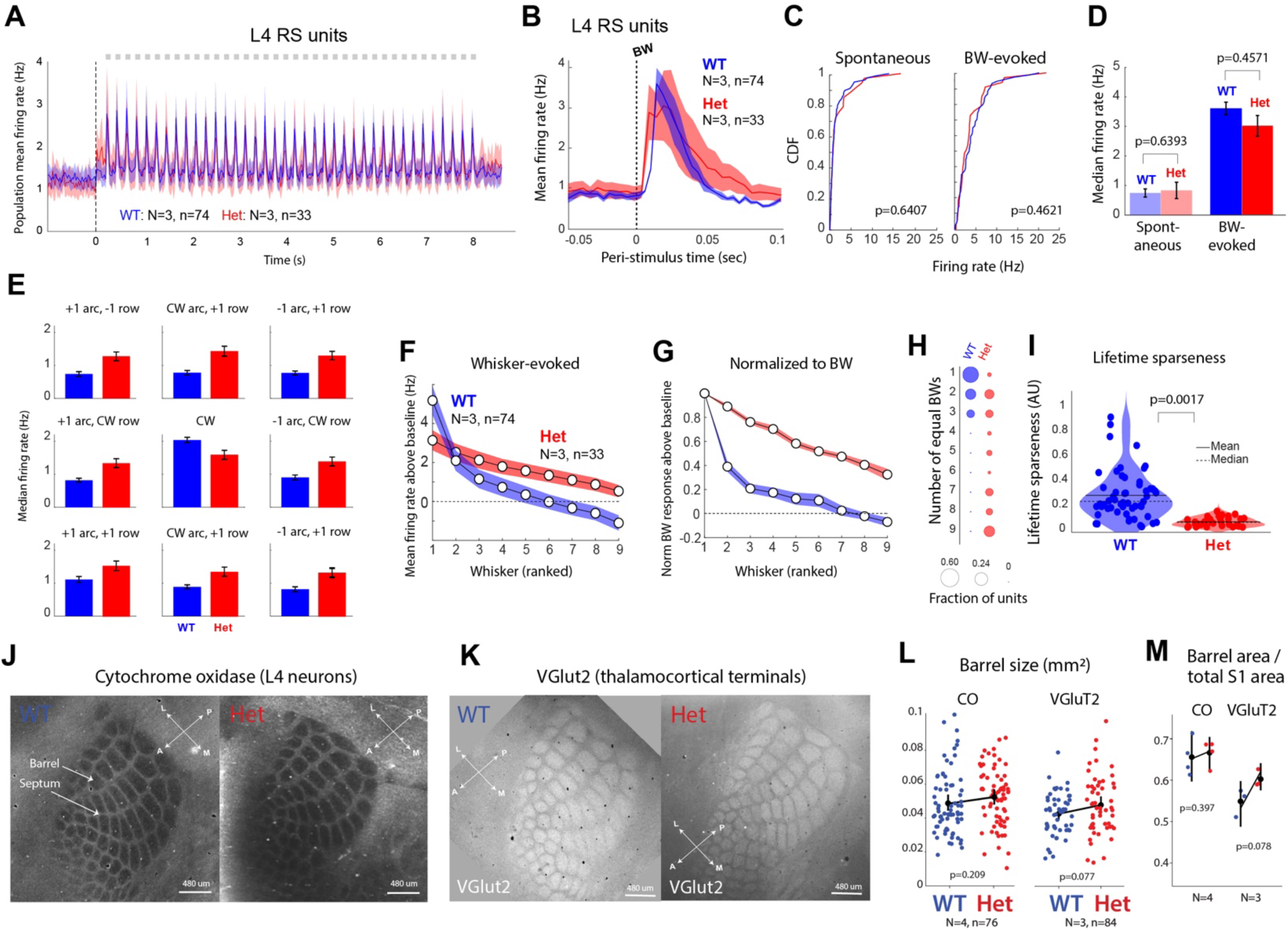
Broad somatotopic tuning in L4 with normal macroscopic thalamocortical patterning. **A)** Mean population firing for L4 RS units across the average trial. Squares are whisker stimuli. WT: N=3 mice, 74 units. Het: N=3 mice, 33 units. **B)** Mean BW-evoked PSTH for WT and Het units. Bars are SEM. **C)** Cumulative density function for spontaneous and BW-evoked firing across units. **D)** Bootstrapped median spontaneous and BW-evoked firing across genotypes. Error bars are 68% CI. P-values from permutation tests. **E)** Mean somatotopic tuning curve for L4 RS units by genotype, showing broader tuning for Het units. N’s. **F-G)**. Mean rank-ordered whisker tuning curves and normalized tuning curves for the same units as in A-E. **H)** Distribution of number of equal BWs. **I)** Lifetime sparseness for each unit. **J**) Cytochrome oxidase-stained section through L4 for an example WT and Het mouse. **K)** Same, but for thalamocortical afferents labeled by anti-VGluT2 staining. **L)** Barrel area from CO-stained and VGluT2-stained sections. Black dot=mean, vertical black line=95% CI. Unpaired t-test, alpha=0.05. N=mice, n=barrels. **M**) Assessment of barrel column vs. septa area, measured as fraction of total barrel field area that is occupied by barrel columns. Black dot=mean. Unpaired t-test, alpha=0.05. N=mice (1 barrel field per mouse).

**Figure 5:**
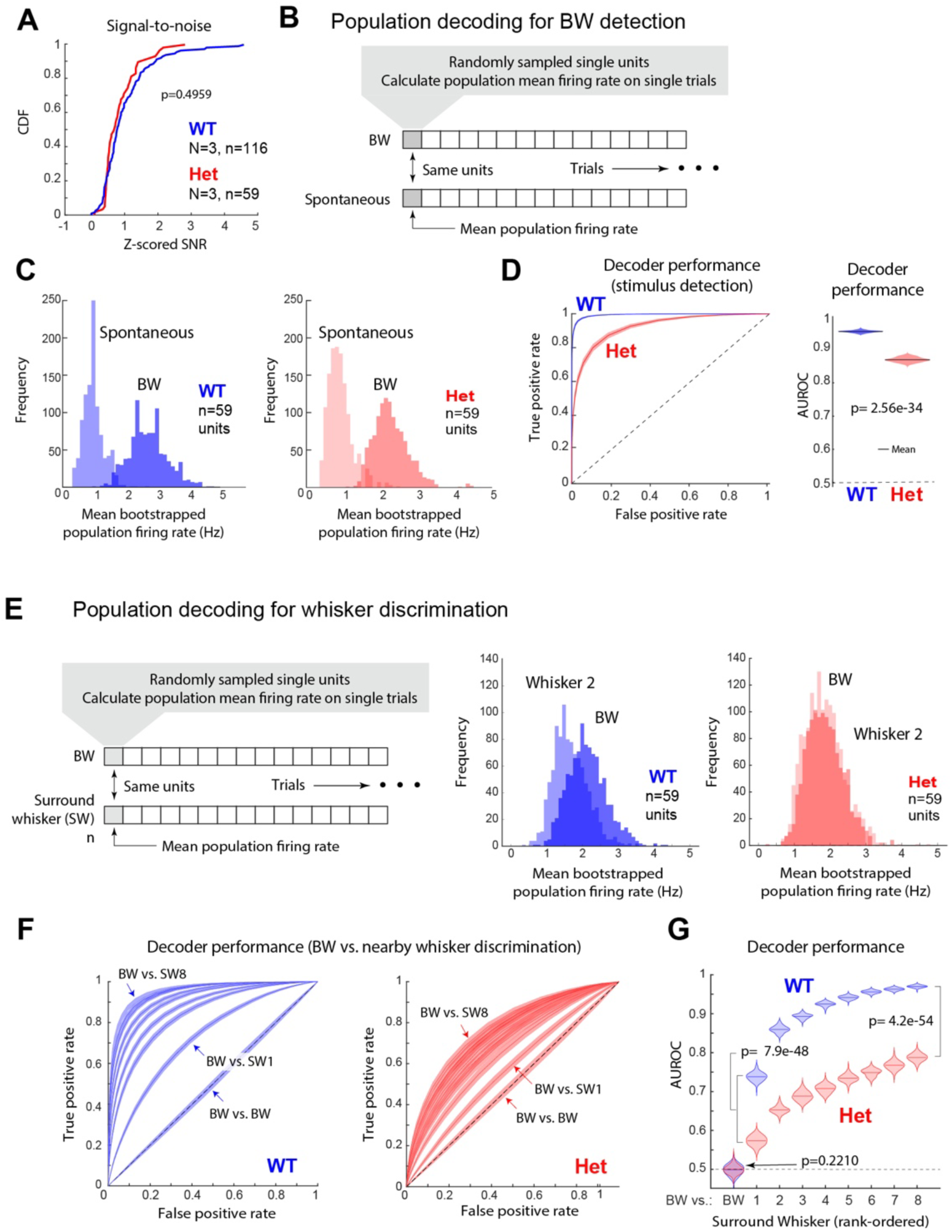
Preferential impairment in neural decoding for discrimination, rather than detection, in *Scn2a^+/−^* mice. **A)** *Z*-scored signal-to-noise ratio for BW-evoked responses in L2/3 RS units. Statistics are from two-sample KS test. **B-D**) Decoding of BW stimulus presence from L2/3 RS population spiking on single trials. C: 1000 bootstrap iterations of population mean firing rate for 59 units sampled from 3 WT mice or 3 Scn2aHet mice. D, left: ROC curves for discriminating BW from blank stimuli based on population firing rate. Error shading is bootstrapped 68% CI across 1000 runs of the decoder. D, right: Distribution of AUROC, mean (solid line), and median (dashed line) across 1000 runs of the decoder. Chance performance is 0.5. Statistics are from permutation test. **E-G**) Decoding of BW stimulus identity from itself or 8 other whiskers from L2/3 RS population spiking on single trials. Same method and conventions as B-D.

To test whether the development of basic thalamocortical projection topography was disrupted in *Scn2a+/−* mice, we used cytochrome oxidase (CO) to label postsynaptic L4 neurons, and vGluT2 immunostaining to label thalamocortical axon terminals, at P22. Postsynaptic barrels were anatomically normal in *Scn2a+/−* (mean barrel size [mm^2^]: WT 0.046 ± 0.002, *Scn2a^+/−^* 0.053 ± 0.002; p=0.20, t-test), as was the fraction of the total barrel field containing barrels (mean: WT 0.66, *Scn2a^+/−^*0.67; p=0.3967, t-test) (**Fig. 5J, L-M**), and relative CO intensity in barrels vs. septa (not shown). Presynaptic thalamocortical patterning was also normal (mean VGluT2 barrel size [mm^2^]: WT 0.042 ± 0.001, *Scn2a^+/−^* 0.047 ± 0.001; p=0.0774, un-paired t-test), mean barrel fraction: WT 0.57, *Scn2a^+/−^*0.60; p=0.2091, un-paired t-test) (**Fig. 5K, L-M**). Therefore, the dramatic broadening of somatotopic tuning in L2/3 and L4 does not reflect macroscopic disorganization of the thalamocortical projection.

### Neural decoding for tactile detection and discrimination

These findings of broader neural tuning, but no major changes in spontaneous or whisker-evoked firing rate, predict that neural populations in *Scn2a^+/−^*mice will have modest or no impairment in encoding whether a whisker stimulus occurred (detection), but may have strong impairment in distinguishing between different whisker stimuli (discrimination). To test this, we first examined coding for stimulus detection, using two approaches. First, we *z*-scored the BW-evoked firing rate of each L2/3 RS single unit to its spontaneous firing rate (from blank trials), which is a measure of signal-to-noise ratio (SNR) for stimulus detection in single units. There was a slight trend for lower SNR in *Scn2a^+/−^*units, but this was not significant (KS test, p=0.4959) (**Fig. 5A**). Second, we constructed a simple single-trial population decoder that tested how well population mean firing rate of L2/3 RS units could detect BW stimuli relative to blank stimuli on single trials. To do this, we compiled all L2/3 RS units into a single virtual population for each genotype, and used bootstrapping to calculate population average firing rate for a fixed number of sampled units on single BW or blank trials. Using receiver operating characteristic (ROC) analysis, we quantified how these virtual single-trial population firing rates distinguished BW from blank trials. The area under the ROC curve (AUROC) represents the optimal performance of a single trial firing rate-based decoder^64^, and expresses the likelihood that an ideal observer correctly classifies trial type (i.e. BW vs. blank trial) from mean population activity (**Fig. 5B-C**). Detection performance was assessed over 1000 iterations of each decoder to calculate confidence intervals, and was slightly but significantly lower for *Scn2a^+/−^*units (mean AUROC: 0.87) than for WT (0.96; p=2.56e-34, Wilcoxon rank sum) (**Fig. 3D**). We found nearly identical results for population decoding of the CW (mean AUROC, *Scn2a*+/−: 0.85, WT: 0.93, p=5e-21, Wilcoxon rank sum) (not shown). Thus, neurometric stimulus detection was unaffected on a single-unit level, but modestly impaired on the population level (96% vs. 87%) in *Scn2a^+/−^* mice.

Next, we tested for impaired neural population coding for whisker discrimination. Using a similar ROC analysis, we compared how well the mean firing rate of virtual L2/3 RS unit populations could distinguish the BW from other nearby whiskers on single trials (**Fig. 5E-G**). For WT mice, AUROC was at chance (0.5) for the BW vs. BW comparison, 0.74 for BW vs. the strongest surround whisker (SW1), and 0.96 for the BW vs. the weakest surround whisker (SW8). This represents high performance at discriminating between nearby whiskers. For *Scn2a^+/−^* mice, AUROC values were systematically lower, being 0.57 for BW vs. SW1 and 0.78 for BW vs. SW8 (**Fig. 5G**). Thus, the performance deficit in *Scn2a^+/−^*mice for population neurometric discrimination between whiskers (21% worse than WT for BW-SW1, 22% worse for BW-SW8, **Fig. 5G**) was much greater than the deficit for neurometric whisker detection (9% worse than WT, **Fig. 5D**). Thus, single-trial analysis indicates the greatest neural coding impairment is in discrimination between whiskers, consistent with a primary deficit in coding distinct sensory features.

### CRISPRa treatment in young adult *Scn2a+/−* mice rescues degraded tactile coding

For treatment of *Scn2a* loss-of-function disorder and other NDDs, a critical unknown is whether restoration of gene expression can normalize sensory or cognitive phenotypes, and if so whether rescue needs to take place during early development, or could also be effective at older, post-critical period ages. In *Scn2a*+/− mice, because macroscopic projections appear normal (**Fig. 4**), but dendritic excitability and LTP are impaired and synapses appear immature^19,47,65^, we hypothesized that that activity-dependent synapse maturation has failed, so that functional microcircuits remain unrefined at the synapse level. If so, restoration of *Scn2a* in older mice may rescue cellular excitability and activity- dependent synapse maturation, enabling circuits to mature and sensory processing to recover. We therefore tested whether CRISPRa upregulation of *Scn2a* gene expression in young adult mice after early S1 critical periods^66–69^ rescues degraded tactile coding or maps.

We delivered recombinant viral vectors (rAAVs) carrying CRISPRa and a guide RNA to the *Scn2a* promoter^70,71^. This approach, applied as a single dose in young adult *Scn2a+/−* mice (P30-40), increases *Scn2a* mRNA expression by 1.5-fold and rescues cellular and synaptic phenotypes in PYR cells in mPFC^47^. Whether this can rescue cortical circuit- and coding-level deficits is unknown. We injected *Scn2a+/−* and WT mice at P30-40 with two viral vectors: i) rAAV-sadCas9-VP64 (a catalytically inactive sadCas9 coupled to the transcriptional activator VP64) and ii) rAAV-*Scn2a*-sgRNA-mCherry (guide RNA to target the *Scn2a* promoter, plus mCherry). The combination of viruses (i) and (ii), termed *Scn2a*-rAAV-CRISPRa, was injected systemically via the retro-orbital sinus, which upregulates *Scn2a* expression throughout the brain^47,70^. As a control, separate mice were injected with virus (i) plus rAAV-EMPTY-mCherry (termed virus iii), which lacks the *Scn2a* guide RNA^47,70,72^. Experimental groups were termed CRISPRaHet, CRISPRaWT, EmptyHet, and EmptyWT, based on their genotype and whether they received viruses (i) and (ii), or viruses (i) and (iii). All mice received a single dose. Critical periods in L2/3 of S1 end by ∼P16, and CRISPRa-mediated gene upregulation requires several weeks with this strategy, so it is likely to drive functionally meaningful expression at ∼P50-60 (**Fig. 6A**). Only cells with endogenous *Scn2a* gene expression should undergo CRISPRa-induced upregulation with this approach^46^.

**Figure 6:**
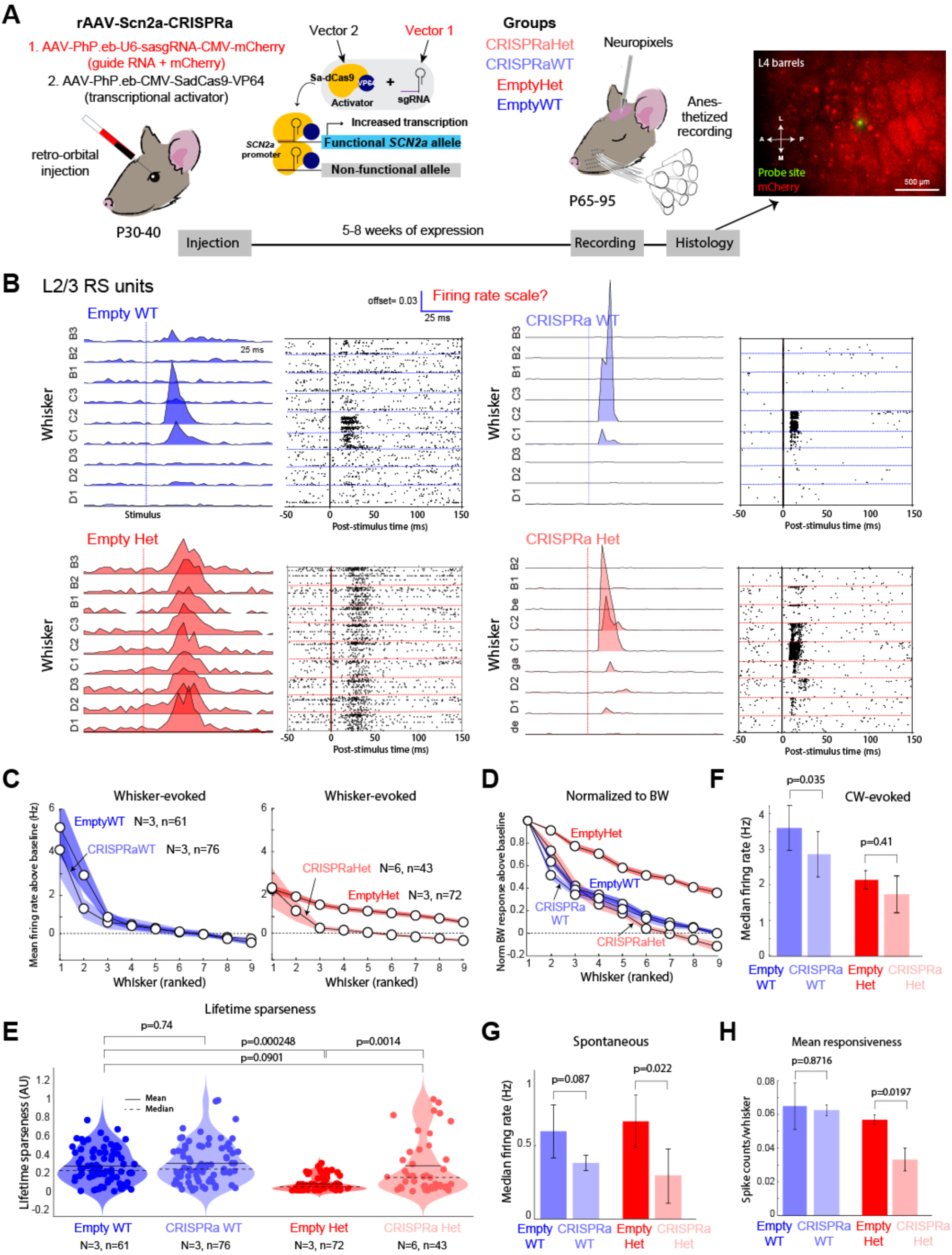
CRISPRa rescues degraded tactile coding in young adult *Scn2a^+/−^* mice. **A**) Design of CRISPRa experiments in mice using systemic delivery of CRISPRa via retro-orbital injections. Right image: mCherry expression (red) and NeuroPixels probe penetration site (green) in a CRISPRa-injected mouse. Section is from L4. **B**) Representative 9 whisker somatotopic receptive fields in L2/3 RS units from EmptyWT, EmptyHet, CRISPRaWT, and CRISPRaHet mice. CRISPRa denotes mice injected with full dual-vector strategy in A. Empty denotes mice injected with one vector lacking the Scn2a guide RNA in A. Vertical line: whisker deflection. Raster and PSTH are for the same unit. **C**) Mean rank-ordered and baseline-subtracted whisker tuning curves. Conventions as in Fig. 3B. All error bars are 68% CI. **D**) Rank-ordered and baseline subtracted whisker tuning curves normalized to BW response. Conventions as in Fig. 3C. **E**) Lifetime sparseness. Statistics are from Tukey-Kramer tests. Alpha=0.05. N=mice, n=units. **F**) Bootstrapped median CW-evoked firing rates. Statistics are from Wilcoxon rank-sum tests and alpha=0.025 for multiple comparisons in F-H. **G**) Bootstrapped median spontaneous firing rates.

We performed *in vivo* extracellular single-unit recordings with Neuropixels probes and recorded ISOI maps under light anesthesia, 5–8 weeks after the injection. Spike recordings spanned L2-L5b. Only mice with significant mCherry expression in wS1 were included in analysis (see Methods). We found that CRISPRaHet mice had larger extracellular spike amplitude for L2/3 and L5 RS units than EmptyHet mice, consistent with successful *Scn2a* upregulation in PYR cells (**Fig. S6A**).

L2/3 RS units in EmptyHet mice exhibited much broader whisker somatotopic tuning than in EmptyWT (median lifetime sparseness: EmptyWT 0.25, EmptyHet 0.08; Tukey-Kramer test, p=2.5e-4) (**Fig**. **6B-E**). This confirms that *Scn2a*+/− mice show broad single-unit whisker receptive fields under anesthesia, as in awake mice. Whisker tuning was much sharper in CRISPRaHet mice for L2/3 RS units compared to EmptyHet mice, evident in individual examples, mean rank-ordered tuning curves, and lifetime sparseness (**Fig. 6B-E**). Lifetime sparseness was higher (sharper tuning) in CRISPRaHet mice compared to EmptyHet (median: CRISPRaHet 0.18, EmptyHet 0.08; Tukey-Kramer test, p=0.0014), and was not different from EmptyWT (median: 0.25; p=0.0901 relative to CRISPRaHet) (**Fig. 6E**). Thus, CRISPRa treatment rescued single-unit somatotopic tuning in *Scn2a+/−* mice. By contrast, CRISPRa in WT mice did not substantially affect somatotopic tuning (**Fig. 6B-D**) or lifetime sparseness (median: CRISPRaWT 0.27, EmptyWT 0.25, p=0.74) (**Fig. 6E**). CRISPRa rescue of tuning sharpness in *Scn2a+/−* mice was also apparent for receptive field center tuning sharpness (**Fig. S6B**), and for every mouse when examined individually (**Fig. S7A**).

Despite sharpening tuning, *Scn2a*-rAAV-CRISPRa treatment did not elevate CW-evoked firing rates in L2/3 RS units. If anything, there was a non-significant trend for lower CW-evoked firing rates in CRISPRaHet and CRISPRaWT, relative to EmptyHet and EmptyWT (bootstrapped median firing rate: EmptyWT 3.63 ± 0.6 Hz, CRISPRaWT 2.84 ± 0.54, permutation test, p=0.035, EmptyHet 2.29 ± 0.23, CRISPRaHet 1.89 ± 0.42, p=0.41) (**Fig. 6F**). Spontaneous firing rates were also modestly reduced (**Fig. 6G**). Though CW-evoked firing rate was unchanged, CRISPRa did reduce mean responsiveness to all 9 whiskers in *Scn2a*Het mice (mean: EmptyHet 0.058 ± 0.007 spikes/whisker, CRISPRaHet 0.033 ± 0.01, permutation test, p=0.0197), which reflects reduced SW-evoked spiking (**Fig. 6H).** Thus, CRISPRa treatment sharpened tuning curves in *Scn2a*^+/−^ mice by reducing SW-evoked firing rate, while generally preserving CW-evoked firing rate. Scn2a-rAAV-CRISPRa treatment also substantially restored sharp tuning for L4 RS units in *Scn2a+/−* mice, assessed with the same analyses as for L2/3 RS units (**Fig. S8**).

To test for rescue of whisker map somatotopy, we performed ISOI in each mouse in these four experimental groups and compared the area of whisker-evoked activation to C1, C2, and C3 whisker deflection (**Fig. 7**). EmptyHet mice had much larger C1 activation areas than EmptyWT mice (Friedman’s test, p=0.0133), recapitulating the genotype effect seen in noninjected mice (**Fig. 7A-B**). C1 activation areas in EmptyWT mice resembled those in non-injected WT mice, and EmptyHet mice resembled those in non-injected *Scn2a+/−* mice, suggesting that viral injection, and CRISPRa and *mCherry* expression, did not alter somatotopy or perturb the ISOI signal (**Fig. 7B**). Strikingly, CRISPRaHet mice showed much smaller, more focused C1 ISOI activation area than EmptyHet mice (Friedman’s test, p=0.0008) or non-injected *Scn2a+/−* mice, which was observable in 5/6 CRISPRaHet mice (**Fig. 7B**). These findings were also true for the C2 and C3 whiskers (**Fig. 7B**). Thus, CRISPRa rescued the sharp whisker map, which occurred without global changes in signal intensity (**Table S2**).

**Figure 7:**
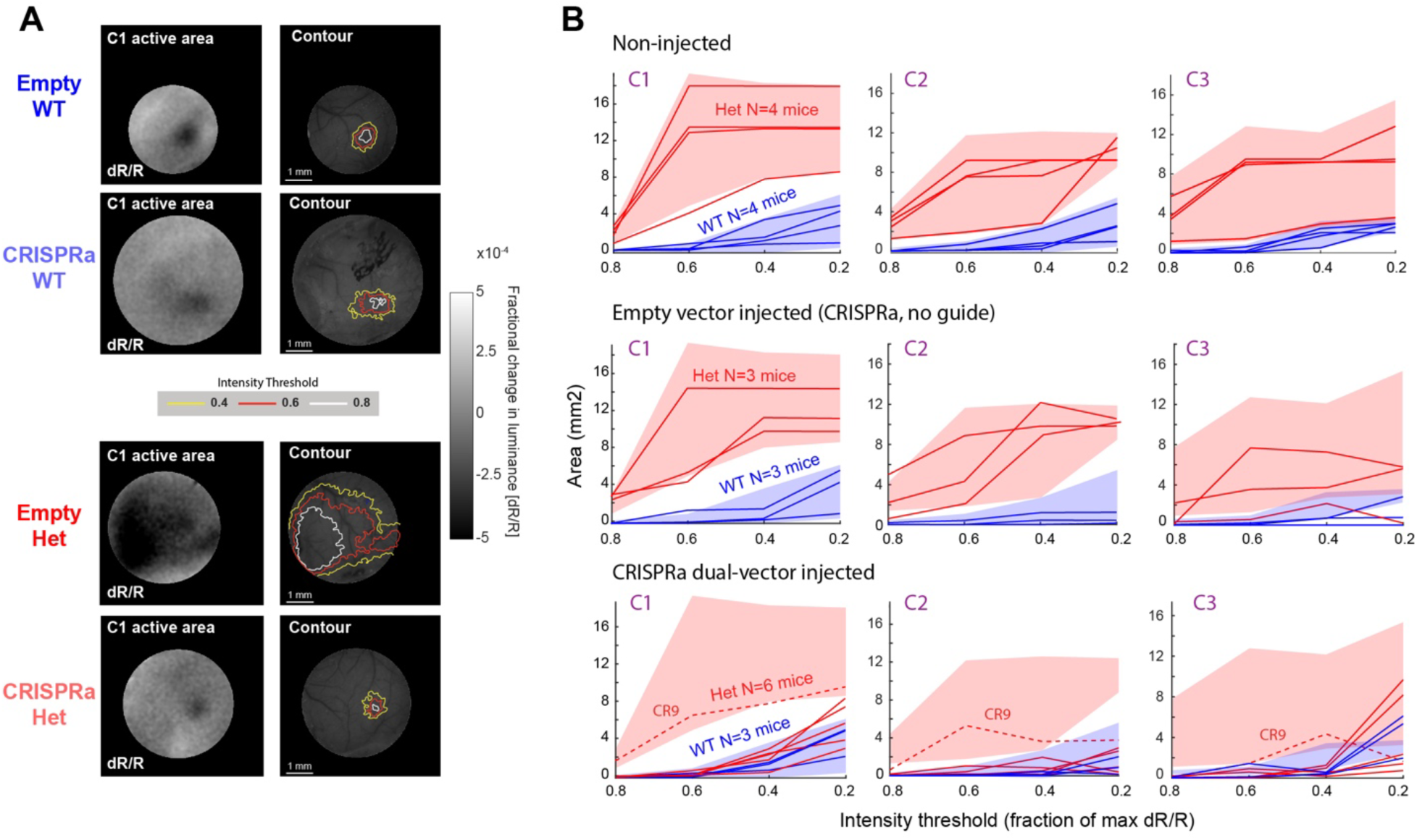
CRISPRa rescue of degraded map somatotopy in young adult *Scn2a^+/−^* mice. **A**) Representative whisker active areas (dR/R: change in reflectance relative to pre-stimulus baseline) and overlaid contours for three intensity thresholds (fraction of maximum dR/R) over mouse skull and vasculature. Yellow: 40% of max dR/R, orange: 60%, white: 80%. CRISPRa: mice injected with full dual-vector strategy. Empty: mice injected with one vector lacking the Scn2a guide RNA. **B**) Quantification of C1, C2, or C3-evoked whisker activation area for wild-type (blue) or Scn2aHet (red) mice (individual lines), in different CRISPRa injection conditions. Top, un-injected mice (data are reproduced from Fig. 4C). Middle, mice injected with EMPTY-rAAV-CRISPRa. Bottom, mice injected with Scn2a-rAAV-CRISPRa. Each line is one mouse. Shaded regions are 95% CI for the non-injected mice (top panel). Dashed line indicates mouse CR9.

In a separate control experiment, we confirmed that injection of the CRISPRa vector, virus (i), without either virus (ii) or (iii), did not by itself alter whisker tuning of L2/3 RS units. WT mice injected with virus (i) alone (N=3 mice) showed sharp whisker tuning for L2/3 RS units, indistinguishable from EMPTY WT mice, and *Scn2a+/−* mice injected with virus (i) alone (N=3 mice) had broad whisker tuning, indistinguishable from EMPTYHet mice. This confirms that the sadCas9-VP64 vector does not produce measurable changes in tuning without a guide RNA to *Scn2a* (**Fig. S9**).

### Consistency of CRISPRa rescue across individual mice

We analyzed rescue of single-unit L2/3 RS tuning sharpness separately for each mouse (**Fig. S7**). All EmptyHet mice showed broader tuning than EmptyWT mice (median lifetime sparseness, Het1: 0.0212, Het2: 0.0104, Het3: 0.0329; combined EmptyWT: 0.0733, permutation test for median < combined EmptyWT: p=0.002, p=0.004, p=0.046, these are adjusted p-values after FDR correction) (**Fig. S7A**). Of the 6 CRISPRaHet mice, all 6 had sharper tuning than EmptyHets, indicating significant rescue (median lifetime sparseness, CR2: 0.1142, CR5: 0.3397, CR6: 0.0828, CR7: 0.0424, CR8: 0.0892, CR9: 0.0475; combined EmptyHet mice: 0.0211, permutation test for median > combined EmptyHet, p=1e-3, 3e-13, 3e-3,1.3e-4,1.5e-6,7e-3, after FDR correction) (**Fig. S7A**). To test whether rescue was complete, we performed equivalence tests asking whether each mouse’s median lifetime sparseness was equivalent to (within a possible 33% effect size)^73^ the combined EmptyWT distribution. All 6 CRISPRaHet mice were equivalent to EmptyWT by this measure (p=2e-6, 2.2e-9, 3.8e-4, 2.2e-4, 04.7e-3, 2.1e-4, after FDR correction). However, one CRISPRaHet mouse (CR9) had the weakest rescue of single-unit lifetime sparseness, and this was the same mouse that retained broad ISOI activation (**Fig. 7B**; **Fig. S7B**).

We assessed whether rescue of single-unit tuning and map topography correlated with cellular-level rescue of *Scn2a* expression, using mean L2/3 RS extracellular spike amplitude as a physiological proxy of *Scn2a* expression in the recorded column. Across CRISPRaHet mice, rescue of mean L2/3 RS spike amplitude correlated with rescue of single-unit tuning sharpness (**Fig. S7C**). Rescue of ISOI whisker activation area was also related to rescue of spike amplitude, but more loosely (**Fig. S7D**). Mouse CR9 was among a cluster of CRISPRaHet mice with little or no spike amplitude rescue. The other two mice in this cluster, CR6 and CR7, showed more substantial rescue of single-unit tuning sharpness, and complete rescue of ISOI activation area, despite weak or no effect on L2/3 RS spike amplitude. This indicates that substantial rescue of somatotopic tuning and maps can occur even when *Scn2a* expression is increased in only a low fraction of local L2/3 PYR cells.

### PV hypofunction and its rescue in *Scn2a+/−* mice

Hypofunction of PV interneuron circuits occurs in many mouse models of ASD-related NDDs^37,51,74,75^, and was apparent in *Scn2a+/−* mice as a strong reduction in spontaneous and whisker-evoked spiking in L2/3 FS cells (**Fig. 1**). To understand why it occurs in *Scn2a+/−*, we first confirmed that PV cells in S1 do not normally express *Scn2a*, as has been demonstrated for PV cells in mPFC by electrophysiology and from gene and protein expression^17,19,47^. We used RNAscope to probe wS1 sections for expression of mRNA for *Scn2a*, *PV*, and the excitatory cell marker *Slc17a7*. In WT mice, *Scn2a* mRNA was robustly expressed in *Slc17a7+* cells, but *PV*+ neurons did not express *Scn2a* mRNA above background levels. In *Scn2a+/−* mice, *Scn2a* mRNA was reduced in *Slc17a7*+ cells and remained very low in *PV*+ cells (**Fig. 8A-B**). We also did not detect expression of GFP in PV cells in S1 in a knock-in mouse that expresses GFP from the endogenous Scn2a locus, as shown in PFC^47^ (**Fig. S10**). To test functionally whether *Scn2a* contributes to spiking in PV cells, we injected a S5E2 enhancer virus^76^ into S1 to drive tdTomato expression in PV cells in WT or *Scn2a*+/− mice, and made whole-cell recordings in S1 slices at P18-22 from L2/3 PV cells, identified by viral expression and fast-spiking properties. In each cell, we injected current to evoke spiking (**Fig. 8C-D**). Analyzing spikes just above rheobase, we found no difference in mean spike waveform, phase-plane plots, spike slope (dV/dt) or spike amplitude between WT and *Scn2a*+/− mice, indicating that *Scn2a* is not functionally expressed in PV cells in S1, similar to results in PFC^19^ (**Fig. 8E-F**).

**Figure 8:**
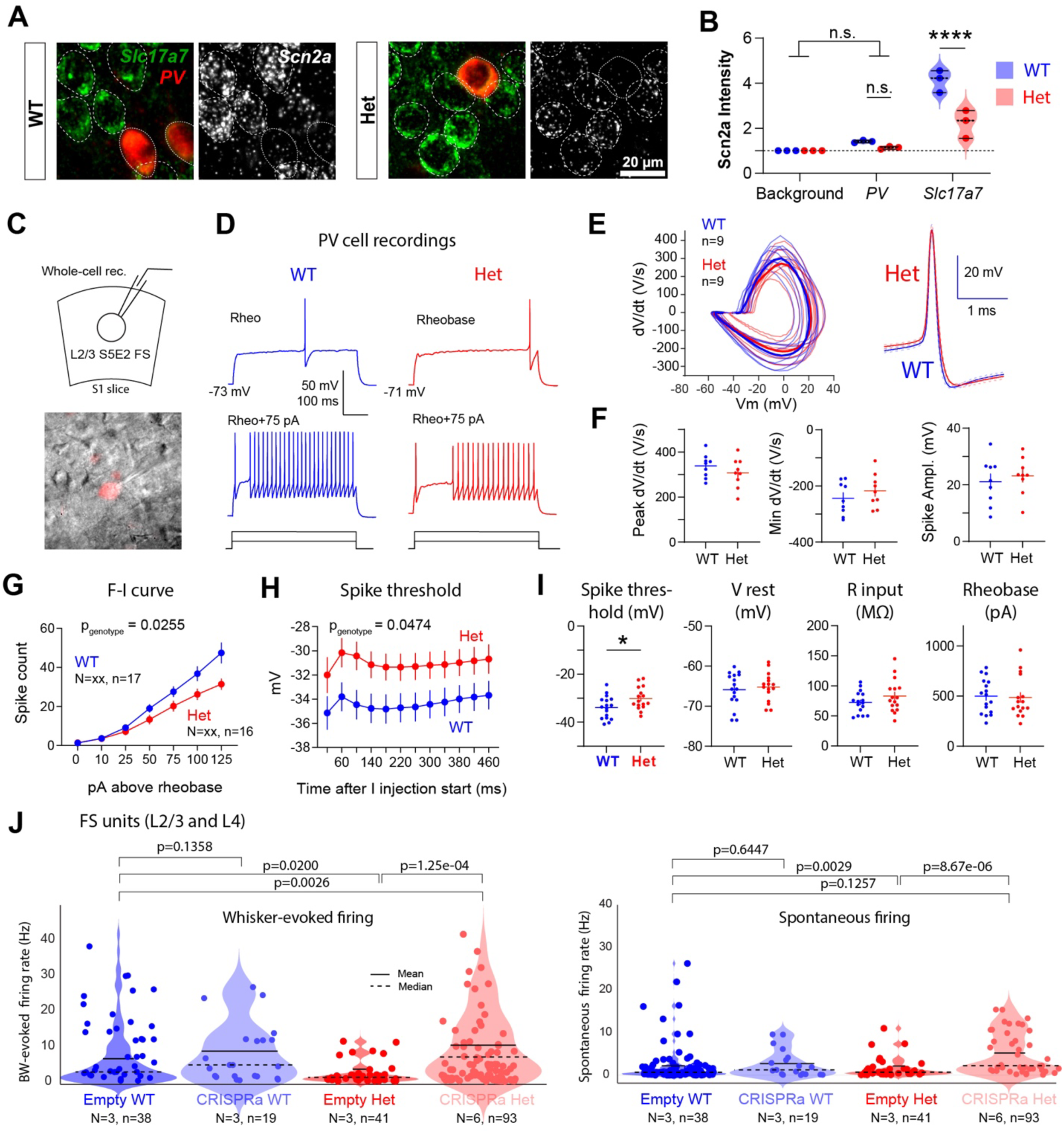
Secondary PV hypofunction in Scn2a+/− mice and rescue by CRISPRa. **A)** Representative RNAscope images showing expression of *Scn2a* mRNA in *PV*+ and *Slc17a7*+ cells in L2/3 of S1. **B)** Quantification of *Scn2a* mRNA in PYR cells (*Slc17a7*+) and PV cells (*PV*+) in WT and Het mice normalized to background signal. Areas of overlap between cells were excluded from analysis. Data are from 3 mice per condition. (Genotype: ***p=0.0005, Cell type: ****p<0.0001, Interaction: ***p=0.0006, Two-way ANOVA with Šídák’s multiple comparisons test). **C)** Whole-cell recordings from PV cells in S1 slices identified using a S5E2 enhancer virus. **D)** Spike trains elicited by current injection in two example L2/3 PV cells. **E-F)** Spike shape in PV cells was identical between WT and *Scn2a*+/− mice, assessed by mean spike waveform, phase-plane plot, spike velocity, and spike amplitude (WT: 3 mice, n = 9 cells; Het: 3 mice, 9 cells; max velocity: p=0.29; min velocity: p=0.35, amplitude: p=0.57, Unpaired t-test). **G)** F-I curve in L2/3 PV cells is reduced in *Scn2a*+/− mice (Ival x Genotype: ***p =0.0003, Genotype: *p = 0.03, Two-way RM ANOVA). **H,** Spike threshold is elevated in *Scn2a*+/− mice (Time Bin X Genotype: *p = 0.02, Genotype: *p = 0.045, Mixed-effects model) **I,** Spike threshold (*p = 0.03), resting potential (Vrest, p = 0.65), input resistance (Rinput, p = 0.21), and rheobase (p = 0.85) across genotypes. Unpaired t-test. Each dot is one cell. For Panels G-I, WT: 6 mice, n = 17 cells; Het: 5 mice, 16 cells. **J**) BW-evoked and spontaneous firing rate measured in vivo for L2/3 C L4 FS units. These units were recorded during the same experiments as the RS units reported in Fig. 1 and Fig. 4 (intermixed at the same recording sites). Statistics are two-sample KS test. Alpha=0.05.

This suggests that reduced spiking in L2/3 FS cells in *Scn2a*+/− mice is not a direct effect of *Scn2a* loss in PV cells, and instead is an indirect, secondary effect of *Scn2a* haploinsufficiency in other cells, presumably PYR cells. S1 PV cells are known to exhibit a powerful form of compensatory plasticity in which they reduce their intrinsic excitability in response to transient reductions in network activity, implementing a network homeostat that stabilizes PYR firing rate^77–79^. This reduction in PV intrinsic excitability is mediated by an increase in PV spike threshold^78,80^. To test whether this mechanism contributes to PV hypoactivity in *Scn2a*+/− mice, we measured F-I curves for PV cells in S1 slices, and found that *Scn2a*+/− PV cells had reduced F-I curves and elevated spike threshold relative to WT, without changes in rheobase, Vrest, or Rinput, suggesting the involvement of this compensatory mechanism (**Fig. 8G-I**).

To test whether CRISPRa rescue of PYR sensory tuning was associated with recovery of PV spiking, we analyzed spontaneous and BW-evoked firing rates in FS units in vivo (combining across L2/3 and L4 to improve sample size). BW-evoked firing in FS units was significantly lower in EmptyHet relative to EmptyWT (Tukey-Kramer test, p=0.0200), as was spontaneous firing in FS units (p=0.0029) (**Fig. 8J**), confirming the findings from L2/3 FS units in awake, non-injected *Scn2a^+/−^* mice (**Fig. 1M-O**). In CRISPRaHet mice, BW-evoked firing rate for FS units was increased above EmptyHet (Tukey-Kramer test, p=1.3e-04), bringing it slightly above Empty WT levels (p=2.6e-3). The same was true for spontaneous firing rate (p=8.7e-06), which was restored to a level indistinguishable from WT (p=0.12) (**Fig. 8K**). Thus, CRISPRa upregulation of *Scn2a* not only recovered L2/3 RS unit spiking and tuning properties, but also recovered FS spike rates.

## DISCUSSION

Neural coding in *Scn2a+/−* mice was dominated by a profound blurring of single-neuron somatotopic tuning and functional map somatotopy, despite largely normal overall PYR cell firing rates (**Figs. 1-4**). Degraded tuning and maps were observed in every mouse, and caused a major deficit in single-trial population decoding of whisker identity, with a smaller deficit in decoding for tactile detection (**Fig. 5**). This is the most profound neural coding impairment in sensory cortex reported for any transgenic mouse model of monogenic ASD (including *Cntnap2−/−*^48,50^, *Ube3am-/p+*^81^, *Fmr1−/−*^48,51^, and *MeCP2−/−*^82^, and *Syngap1*^83^). Only cortical layers L2/3 and L4 were examined, but population activity in L2/3 is closely related to perceptual behavior^84,85^. These findings therefore reveal a major impairment in discriminative sensory processing for touch in wS1, which may be the mouse tactile analog of cortical visual impairment (CVI) which is common in *Scn2a*-related disorders^5,6^.

Despite the fact that *Scn2a* haploinsufficiency directly reduces PYR cell dendritic excitability and weakens excitatory synapses^19^, L2/3 and L4 RS units showed normal spontaneous and whisker-evoked firing rates. Thus, *Scn2a+/−* falls within a phenotypic cluster of ASD model mice with blurred tuning and maps, but largely normal PYR firing rates (*Cntnap2−/−*, *Ube3am-/p+*, *Mecp2−/−*, and many studies in *Fmr1- /y*), predicting a blurry or confusing cortical sensory world^49^. Stable PYR firing rates may reflect known cellular-level compensation to *Scn2a* loss by reduced NaV1.2-dependent activation of dendritic K currents that restores PYR cell excitability^18,86^, or by classical network compensation to reduced PYR firing, including homeostatic synaptic up-scaling of PYR-PYR synapses^87^ or weakening of PV cell intrinsic excitability^77^.

Blurred neural coding may be a direct consequence of weak excitatory synapses onto PYR cells in *Scn2a+/−* mice. *Scn2a+/−* synapses show impaired LTP, low AMPA:NMDA ratio, and a high fraction of silent synapses and immature spines^19^. This suggests a loss of activity-dependent synapse strengthening that may prevent experience-dependent synaptic refinement during early critical periods, yielding blurred functional microcircuits and broad tuning and maps. Normal macroscopic thalamocortical topography in L4 (**Fig. 4**) suggests that coarse circuit patterning is intact, implicating failed refinement at the synaptic level in tuning loss. Another contributor to blurred coding could be reduced activity in PV interneurons, which was a surprising finding *in vivo* (**Fig. 1**) and involves a reduction in PV intrinsic excitability (**Fig. 8**). Cortical PV cells are not thought to express *Scn2a*, consistent with our mRNA expression and electrophysiology data (**Fig. 8**). This suggests that PV cell hypofunction reflects network- level compensation to *Scn2a* loss in PYR cells or other neurons. PV cells in S1 are known to reduce their intrinsic excitability in response to transient reductions in network activity, thus stabilizing PYR firing rate^78,79^. The resulting drop in PV activity may blur PYR tuning and maps by impairing developmental critical periods^88^, or by contributing to a general loss of inhibition that may acutely broaden PYR sensory tuning^89^. We cannot rule out that some PV cells may modestly express NaV1.2 protein, causing a direct reduction in PV excitability in *Scn2a*+/− mice. Substantial changes in gene expression in MGE-derived interneurons were recently found in mPFC of a *Scn2a* knockdown mouse model, supporting the idea that, whether directly or indirectly, inhibitory circuits can be a locus of abnormal cortical function in *Scn2a*-related disorders^90^.

We found that CRISPRa-mediated transcriptional activation of *Scn2a* (rAAV-Scn2a-CRISPRa), applied in young adult mice several weeks after the closure of S1 critical periods, robustly rescued neural coding phenotypes in *Scn2a+/−* mice. Subcortical and thalamocortical circuits in wS1 develop and refine by ∼P5, intracolumnar and cross-columnar excitatory circuits by ∼P16, and inhibitory circuits by ∼P25^91^. After Scn2a-rAAV-CRISPRa injection at P30-40, which is estimated to upregulate *Scn2a* gene expression by ∼P50-60, mean whisker tuning for L2/3 RS units in *Scn2a+/−* mice sharpened to become indistinguishable from wild types (**Fig. 6**), and L4 RS tuning sharpened to nearly this level (**Fig. S8**). Every *Scn2a+/−*mouse showed rescue of L2/3 RS unit tuning (**Fig. S7**), and 5/6 mice showed rescue of whisker map topography by ISOI (**Fig 7**). A physiological proxy for NaV1.2 functional expression in L2/3 PYR cells, the mean L2/3 RS extracellular spike amplitude, was increased by rAAV-Scn2a-CRISPRa, and the extent of increase was correlated with rescue of neural tuning. However, even mice with no detectable increase in RS spike amplitude showed significant rescue of single-unit tuning and map topography, suggesting that blurred tuning and maps can be effectively rescued by *Scn2a* upregulation in a low fraction of local PYR cells (**Fig. S7**). Thus, even though *Scn2a+/−* neural coding phenotypes are severe, they can be rescued in post- critical period animals.

The mechanism of coding rescue by rAAV-Scn2a-CRISPRa is unclear. rAAV-Scn2a-CRISPRa treatment in young adults rescues PYR dendritic excitability and restores the ability of PYR excitatory synapses to undergo LTP^47,71^. This could allow ongoing tactile experience to strengthen excitatory synapses onto PYR cells, effectively opening a delayed critical period that sharpens tuning and maps. rAAV-Scn2a-CRISPRa could also transiently elevate PYR firing rates to drive network homeostasis including upregulation of PV and other inhibitory neuron function to sharpen PYR tuning. We found CRISPRa treatment elevates PV firing rate, suggesting this mechanism plays a role (**Fig. 8**). It is also possible that CRISPRa acts directly in a non-PYR cell type, or subcortically, to sharpen sensory coding in S1.

CRISPRa is a promising therapeutic strategy for genetic disorders caused by heterozygous loss-of- function mutations^72^, which include *Scn2a*-related ASD/ID as well as most syndromic forms of ASD. By directly targeting the endogenous gene regulatory elements, CRISPRa is particularly suited for genes like *Scn2a* with coding sequences too large for direct AAV-mediated gene replacement^72^. A previous study showed that rAAV-CRISPRa-Scn2a, applied in young adults, increases *Scn2a* expression and rescues cellular-level dendritic excitability and synaptic phenotypes in mPFC PYR cells in *Scn2+/−* mice^47^. Our data shows that adult CRISPRa treatment can rescue circuit- and coding-level phenotypes in NDDs, indicating that cortical information processing can be reversed even after early critical periods for circuit development. Thus, CRISPRa therapy or other strategies for increasing NaV1.2 function could be successful in older children or adults.

Together, these findings show a profound impairment in cortical sensory processing in *Scn2a+/−* mice, and that functional restoration of *Scn2a* in young adult, post-critical period mice is effective in rescuing major aspects of information processing in sensory cortex.

## METHODS

### Mouse strains and conditions

Procedures were approved by the UC Berkeley Animal Care and Use Committee, and followed NIH guidelines. *Scn2a*^+/−^ mice were generated by breeding *Scn2a*^+/−^ males (from ref. ^92^) with C57BL6/J females. Pups were genotyped and separated after weaning into cohorts of five or fewer littermates, until used for slice physiology or craniotomy surgery. Post-surgery mice were housed singly. Mice were maintained on a 12/12 light-dark cycle at 20–26°C with 30–70% humidity. Mice of either sex were used in experiments.

### Ex Vivo Electrophysiology

#### Slice Preparation

Postnatal day (P) 18-22 mice were anesthetized with isoflurane and decapitated. Brain slices were prepared using a Leica VT1200S vibratome in chilled, oxygenated, low-sodium, low-calcium Ringer’s solution (in mM: 85 NaCl, 75 sucrose, 25 D-(+)-glucose, 4 MgSO_4_, 2.5 KCl, 1.25 Na_2_HPO_4_, 0.5 ascorbic acid, 25 NaHCO_3_, and 0.5 CaCl_2_, 320 mOsm). Cortical slices (350 µm) were cut from the left hemisphere in the “across-row” plane, oriented 50° toward coronal from the midsagittal plane and 35° from vertical. Using this plane, each slice contains one column from each whisker row A–E, and within-column circuits are largely preserved^93,94^. Slices were transferred to standard Ringer’s solution (in mM: 119 NaCl, 26.2 NaHCO_3_, 11 D-(+)-glucose, 1.3 MgSO_4_, 2.5 KCl, 1 NaH_2_PO_4_, 2.5 CaCl_2_, 300 mOsm) for 30 min at 30°C and kept at room temperature for 0.5–6 h until recording.

#### Whole cell recordings in S1 slices

Recordings were made at 30°C in standard Ringer’s solution. Barrel columns and layers were identified by transillumination. For L2/3 PYR cells, visually guided patching was performed using infrared differential interference contrast optics at 40X, with PYR cells identified by soma shape. PYR cells were located 100 –240 µm below the L1–L2 boundary, within 100 µm tangentially of column center. Recordings were targeted to the C, D and E columns. Whole-cell recording was performed with 3–5 MΩ pipettes using a Multiclamp 700B Amplifier (Molecular Devices). For measurements of action potential waveform, data were acquired at 50 kHz and filtered at 20 kHz. For all other measurements, data were acquired at 10-20 kHz and filtered at 3-10 kHz.

For current-clamp recordings, pipette capacitance was compensated by 50% of the fast capacitance measured under gigaohm seal conditions in voltage-clamp prior to establishing a whole-cell configuration, and the bridge was balanced. Current-clamp recordings were performed using K gluconate internal solution (mM: 116 K gluconate, 20 HEPES, 6 KCl, 2 NaCl, 0.5 EGTA, 4 MgATP, 0.3 NaGTP, and 5 Na_2_-phosphocreatine, pH 7.2, 290 mOsm). Series resistance artifacts were corrected by bridge balance. AP threshold and peak dV/dt measurements were determined from the first AP evoked by a near-rheobase current in pyramidal cells (500 ms duration). Single action potentials from 5 sweeps per cell were averaged to determine AP threshold and peak dV/dt.

For voltage-clamp recordings of mEPSCs, we used cesium gluconate internal solution (in mM: 108 D-gluconic acid, 108 CsOH, 20 HEPES, 5 tetraethylammonium (TEA)-Cl, 2.8 NaCl, 0.4 EGTA, 4 MgATP, 0.3 NaGTP, 5 BAPTA, and 5 QX-314 bromide, pH 7.2, 295 mOsm). The holding potential (Vhold) was corrected for the liquid junction potential (12 mV). Series resistance (Rseries) was monitored in each sweep. Rseries compensation was not used for mEPSC recordings. Cells with series > 15 MΩ were excluded. Cells whose input resistance (Rinput) changed >30% throughout recording were also excluded from analysis. mEPSCs were recorded in voltage clamp at −72mV with 50 µM D-AP5 and 0.1 µM tetrodotoxin (TTX) in the bath. Events were detected using a deconvolution algorithm^95^ with a noise threshold of 3.5x. The detection threshold was 5 pA. A minimum of 300 events were analyzed per cell.

For PV cell recordings, mice (P3) were injected with AAV-S5E2-dTom-nls(dTom) (Vector BioLabs) into S1 cortex using a Nanoject. 414 nl were injected at one site in the left hemisphere (coordinates from lambda: AP: +1.60 mm, ML: +1.75 mm, ∼0.3 mm DV). At P18-22, S1 slices were prepared as described above. Whole-cell current clamp recordings were made as described above, targeted to tdTom fluorescent neurons in L2/3 of C, D, or E columns. Only neurons that exhibited fast-spiking spiking patterns in response to 500-ms current injection were used for data collection.

### RNAscope measurement of mRNA expression

P21 mice were deeply anesthetized with isofluorane and transcardially perfused with 1X RNase-free PBS. Brains were fixed in 4% RNase-free paraformaldehyde (PFA) in PBS (pH 7.4), then allowed to sink in 30% sucrose at 4°C. Across-row slices were made on a sliding microtome at 25 µm. Slices were mounted in PBS and allowed to dry thoroughly on the slide, then post-fixed in 4% RNAse-free PFA for 1 hour at 4°C, followed by serial ethanol dehydration.

Multiplex fluorescence in situ hybridization followed the protocol for ACDBio’s RNAscope Multiplex Fluorescent V2 Assay (Advanced Cell Diagnostics, cat# 323110). Thawed sections underwent H_2_O_2_ permeabilization, 5-minute target retrieval, and Protease III treatment. RNAscope probes for *Slc17a7* (catalog #501101), *PV* (#421931-C4) and *Scn2a (*#887241-C3) were applied and amplified in sequence with TSA Vivid and Opal Polaris dyes (Advanced Cell Diagnostics, cat # 323271, 323272, 323273; Akoya Biosciences, cat # FP1501001KT). Cellular nuclei were counterstained with 1 µg/ml DAPI and mounted with Prolong Glass Antifade Mountant (Thermo Fisher Scientific, cat# P36982). RNAscope runs were performed with both genotypes side-by-side and positive and negative control samples, to control for inter-batch variability. Imaging was performed on a Keyence BZ-X1000 fluorescence microscope, at 60X magnification using optical sectioning in the *Slc17a7* and *Scn2a* channels to improve resolution. Images were acquired in the C, D, or E barrel of L2/3 of S1 and analyzed in FIJI. Z-stacks were max-projected and manual segmentation of cells in the *Slc17a7* (putative excitatory cells) and *PV* channels was performed. Mean intensity per cell was quantified in the Scn2a channel and cells were averaged per mouse. Areas of overlap between cells were excluded from analysis. Three mice per genotype were used across 2 separate technical replicates for reproducibility.

### Headpost implant surgery

Mice used in in vivo recording or intrinsic signal optical imaging experiments underwent headpost implantation. At 2.5–3 months of age, mice were anesthetized with isoflurane (1-2% in O_2_) and administered dexamethasone (2 mg/kg), enrofloxacin (5 mg/kg), and meloxicam (10 mg/kg). A stainless-steel head post was affixed to the skull using cyanoacrylate glue and dental cement. The C1–3 whisker columns in wS1 were localized using transcranial intrinsic signal optical imaging (see below). After imaging, mice recovered on a warming pad and given subcutaneous buprenorphine (0.05 mg/kg) to relieve post-operative pain. All mice with headpost implants were singly-housed, and each cage had a running wheel (Mouse Igloo #K3327, Bio-Serv).

### Sensory behavioral training for awake in vivo recordings

Mice were trained on a head-fixed Go-NoGo visual sensory detection task, to ensure consistent behavioral state for in vivo recordings of whisker sensory tuning. Training began one week after recovery from head post implant surgery. Training sessions took place during the dark (active) phase of the light cycle. Mice were trained 5 days per week.

#### Behavioral apparatus

Mice were transiently anesthetized with isoflurane (4% induction, 1.5% maintenance in O_2_), placed in the behavioral apparatus, and head fixed via the implanted head-post. Eye ointment (product) was applied, and 9 whiskers were inserted into piezo actuators. A solenoid-gated drink port with an infrared (IR) lick sensor was placed within reach of the tongue. An infrared camera monitored the mouse’s movements. A blue LED was mounted in front of the mouse for visual stimulation in the visual detection task. A heating pad maintained body temperature at 37° C. Once the mouse was positioned, anesthesia was discontinued, the heating pad was switched off, and mice were given 10-15 minutes for recovery before behavioral trials began. Behavioral training was conducted in total visual darkness except for the blue LED, with masking noise (continuous white noise, 70 dB) applied to mask sounds from piezo deflection and drink port opening.

#### Trial Structure

Each trial lasted 9.5 s and consisted of a 0.2 s pre-stimulus period, 7.8 s whisker stimulus period, and 1.5 s response window. During the whisker stimulus period, a train of 38 single-whisker stimuli, on randomly chosen whiskers including blanks (no-whisker deflection) was applied with 0.2 s inter-stimulus interval (ISI). A 1.5 sec response window began at the end of the whisker stimulus period. Trials were separated by 3 ± 2 s inter-trial interval (ITI). On Go trials, a blue LED was flashed at the beginning of the response window, cueing the mouse to lick. NoGo trials contained no LED flash. The fraction of Go trials was 70-80%, adjusted daily to maximize the total number of trials and task performance. A lick response was defined as 2 licks within 0.3 sec during the response window. Hit trials (i.e., Go trials with a lick response) were rewarded with 4 µL water immediately following the lick response. False alarms (i.e., NoGo trials with a lick response) were not rewarded. Mice self-initiated each trial by suppressing licking during the ITI. Trials were aborted if mice gave a lick response during the pre- stimulus or whisker stimulus period.

#### Whisker Stimulation

For each recording site, the columnar whisker (CW) and the 8 adjacent surround whiskers (SWs) on the right side of the face were inserted in a 3 × 3 array. Whiskers were not trimmed, but were threaded into a glass capillary tube attached to the piezo, and held in place by rubber cement. Custom behavioral software (written in Igor Pro) delivered whisker deflections. Each whisker deflection was a ramp-hold-return movement (3 deg. deflection, rostrocaudal direction, 5 ms rise – 1 ms hold– 5 ms fall, applied 5 mm from the face). Each stimulus was a single whisker deflection or a blank. Each trial contained a train of 38 whisker stimuli or blanks (with the whisker identity for each whisker deflection chosen pseudorandomly), at 0.2 sec ISI within the train. Blank stimuli were used to assess baseline firing rate of neurons.

#### Behavior quantification

To assess behavioral task engagement, visual detection performance was quantified by d-prime as:

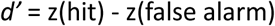

where z = inverse of the normal cumulative distribution function.

#### Training Stages

Mice were water restricted and received water rewards during training, supplemented by additional water to reach a constant volume daily. Daily intake was individually calibrated to maintain an ad lib body weight >80%, and adjusted periodically to maintain body weight and task performance. Training proceeded in stages. In Stage 1, mice were acclimated to handling, the apparatus, and head- fixation. In Stage 2, mice learned to lick for water rewards from the water port. In Stage 3, mice were acclimated to transient anesthesia and recovery on the apparatus. In Stage 4, blue LED flashes were introduced, and mice learned to lick during the response window after LED flashes. This stage did not include whisker stimuli, but trial structure and mean interval of reward availability was matched to the final task structure. At the beginning of this stage, reward was delivered automatically after each LED flash. As the stage progressed, mice were required to lick after the LED flash (during the response window) to obtain reward. When LED-evoked licking was reliable, mice proceeded to Stage 5 in which whisker stimuli were introduced and the whisker stimulus period was gradually lengthened until it was 8.0 seconds. Stage 5 training continued until d-prime >1, HR ≥ 0.8, abort rate < 0.2, and the number of non-aborted trials per day was ≥200.

### Physiological recordings in awake behaving mice

#### *In vivo* recordings in behaving mice

Multiple recording sessions were performed during behavior in each mouse, spaced over 3-5 days, and were complete by P90-250. Recordings were performed blind to genotype. Before the first recording session, the C1, C2, and C3 columns in wS1 were localized by ISOI, and a 0.5-2mm craniotomy was made. During each recording session, the mouse was transiently anesthetized and placed on the behavioral rig as during behavioral training. While the mouse was still under anesthesia, a 32-channel NeuroNexus laminar polytrode (A1×32-Poly2–5mm-50s-177-A16) or a Neuropixels 1.0 probe was inserted radially in wS1, advanced to a tip depth of 500 microns, and allowed to settle for 15-20 minutes while the mouse recovered from anesthesia. A recording ground was secured in the recording chamber. After the mouse recovered, the behavioral and neurophysiological recording session began.

For the awake behaving recordings, 3 mice per genotype were recorded with NeuroNexus probes, and 1 mouse per genotype was recorded with Neuropixels probes. NeuroNexus recording methods are given here. Neuropixels recording methods occurred under the same awake conditions, but followed the recording methods describe for the CRISPRa experiments below.

#### Spike and LFP data acquisition with NeuroNexus probes

For NeuroNexus recordings, signals were sampled (24.4144 kHz) using OpenEx software and stored (TDT RZ5D). For spike detection, signals were bandpass filtered offline (300–6000 Hz) and common average referenced using custom-written MATLAB code. Automatic spike detection was performed within pre-specified tetrode channel groups located within L2/3 or L4, with depth of each channel group defined as the geometric mean of the microdrive depth of the component channels. Layer identity was defined by depth (L2/3: 100 – 413 μm, L4: 414–588 μm below the pia).

Spike sorting for NeuroNexus recordings was performed within each tetrode group using Kilosort2^96^, following by manual curation using Phy (https://github.com/cortex-lab/phy). Isolated clusters were manually inspected for mean spike waveform amplitude, drift stability, and inter-spike interval refractory period violations, which could not exceed 2% for intervals < 1.5 ms. Only well-isolated single units were kept for analysis. Single units were classified as RS or FS based on trough-to-peak distance of the spike waveform at the highest-amplitude recording channel, with a separation criterion of 0.419 ms^48^.

LFP signal was isolated from the same acquired signal as raw spike data, but filtered using a bandpass (10-200 or 300 Hz) and a second-order Butterworth filter. Channels were assigned to layers by depth using the same depth criteria as for single units. For analysis of whisker-evoked LFP in L2/3, the channel in L2/3 with the largest-magnitude negative-going whisker-evoked LFP transient was chosen.

#### Histological determination of recording locations

After the final recording session in each mouse, recording penetration locations were marked with DiI, the animal was sacrificed and the brain fixed in 4% paraformaldehyde. Flattened tangential sections were prepared and stained for cytochrome oxidase (CO) activity as described below. DiI fluorescence and CO staining were imaged, and each marked penetration site was localized relative to barrel column boundaries. Locations of additional unmarked penetrations were interpolated based on microdrive coordinates and surface blood vessels. Recording site locations were compared across mice using a normalized radial coordinate system centered on the barrel centroid, with r = radial position as a fraction of the barrel major axis radius, and theta = angle relative to within-row and within-arc directions, determined from the centroids of neighboring barrels.

### Quantitative analysis of spike and LFP data

#### Analysis of in vivo spike data

Analysis was conducted in MATLAB. Only non-aborted trials (i.e., free from licks during the whisker stimulus period) were analyzed, to avoid contamination by lick-related spiking.

Whisker-evoked and spontaneous firing rate were calculated in the 0-100 ms period following onset of a whisker stimulus or blank stimulus, respectively. To determine whether a given whisker evoked a significant response from a single unit, we performed a permutation test on firing rate between whisker stimulus trials and blank trials. If mean whisker-evoked firing rate was greater than the (1-α) percentile of the null distribution obtained from shufling whisker and blank trial labels, the whisker was concluded to evoke a significant response. Because 9 whiskers were tested for each unit, we used α=0.0055 (α=0.05 / 9 whiskers). A unit was considered whisker responsive if at least one whisker evoked a significant response.

For each whisker-responsive unit, the ‘best whisker’ (BW) was defined as the whisker that evoked the highest firing rate. Any whisker that evoked a response that was statistically indistinguishable from the BW (by permutation test) was considered an equivalent best whisker (eBW). All reported data on BW-evoked responses, number of eBWs, and whisker tuning, are from whisker-responsive units. The *z-*scored signal-to-noise ratio (SNR) for BW responses was calculated for each unit as:

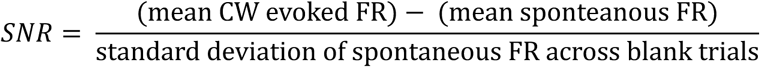

where FR is firing rate.

#### Tuning curves and lifetime sparseness

To calculate a unit’s rank-ordered tuning curve, each whisker stimulus was ranked from highest- to lowest-evoked firing rate for that unit, and the spontaneous firing rate was subtracted. The BW is, by definition, whisker stimulus rank 1. Only units whose BW was the center whisker in the 9-piezo array were included in rank-order tuning analysis, to ensure that all 8 surrounding whiskers (SWs) were sampled. Normalized tuning curves were calculated by dividing each unit’s rank-ordered tuning curve by its BW response. Population rank-order tuning curves were calculated by averaging across the individual rank-ordered tuning curve for each unit.

To quantify tuning selectivity across all 9 whiskers for each unit, we used lifetime sparseness^56^:

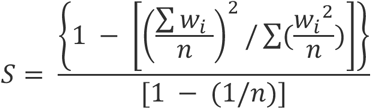

where *w*_*i*_ is the mean whisker-evoked firing rate for the *i*th whisker and *n* is the total number of whiskers. *S* equals zero indicates non-selective tuning tuning, and *S* equals 1 indicates a unit that is perfectly sharp, i.e. responsive to only a single whisker stimulus. To quantify mean responsiveness across all whisker stimuli for each unit, we averaged whisker-evoked spiking response across the 9 whisker stimuli.

#### Population decoding for detection and discrimination

Decoding was performed using receiver-operating characteristic (ROC) analysis of virtual population data, to quantify the ability to discriminate CW-evoked firing rate trials from spontaneous firing rate trials (for detection), or “whisker A” firing rate trials from “whisker B” firing rate trials (for discrimination). For each genotype, a virtual population was compiled by combining units across mice within the genotype, and then randomly subsampling so that the number of units was matched between genotypes. To simulate single trial virtual population activity, one trial was randomly chosen for each unit and mean population firing rate was calculated. From this simulated trial data, ROC analysis was performed to assess discrimination of CW vs. spontaneous trials, or “whisker A” vs. “whisker B” trials, using MATLAB’s perfcurve function. An area-under-the-ROC-curve (AUROC) value of 0.5 indicates chance performance, and 1.0 is perfect performance. Performance was assessed over 1,000 decoder iterations, each using a different random subsampling of units, to calculate confidence intervals for decoding.

#### Local field potential (LFP) analysis

Analysis was performed using custom Python code in Jupyter notebooks. The negative-going LFP peak amplitude was used to quantify each whisker’s response for rank-ordered tuning curves and lifetime sparseness. In addition, we calculated receptive field center tuning sharpness as:

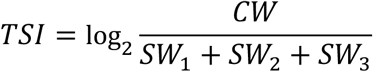

where *CW*, *SW*_1_, *SW*_2_, and *SW*_3_ are the mean peak LFP responses to the CW and the top three rank-ordered surround whiskers, respectively.

### Intrinsic signal optical imaging (ISOI)

#### Image acquisition

Images were acquired using a 12-bit 1M60 Dalsa camera and a macroscope composed of two 50-mm camera lenses^57^. Imaging was performed through the intact skull, using a saline-filled well covered with a glass coverslip. After acquiring a surface vasculature image using green (525 nm) light, the camera was focused down 400-450 μm below the surface blood vessels, illumination was switched to red (630 nm) light, and images (256 pixels × 256 pixels, 6 mm × 6 mm) were acquired at 30–50 fps using custom-written LabView software and binned into 0.5-s bins for analysis. Illumination intensity was adjusted in each experiment to achieve nearly-maximal baseline pixel intensity without any pixel saturation.

On each of 30-40 trials, an ISOI movie was collected while deflecting either the C1, C2 or C3 whisker. Each movie consisted of a 2-sec pre-stimulus baseline period, followed by a train of whisker stimulus pulses to the single whisker (22 pulses, 40 ms inter-pulse interval, each pulse was a ramp-hold-return with 4 ms rise, 1 ms hold, 4 ms fall, 400 µm amplitude rostrocaudal direction).

#### Image analysis

Analysis was performed using custom code in MATLAB. First, a single trial-averaged movie was calculated for each whisker. Whisker responses were analyzed from the 2 frames spanning 1– 1.5 s after whisker stimulus onset, which were averaged before subtracting the mean of 2 baseline frames (−1 to 0 sec) in the same trial. This delta-R image was divided by the mean baseline image to generate a dR/R image, which was then spatially filtered with a 2-dimensional Gaussian (37.5 μm standard deviation). The median of the entire image was subtracted to minimize the effects of slow, global fluctuations in luminance^97^. The strongest pixel was defined as the pixel with the largest negative dR/R value. The WRA for the stimulated whisker was quantified as the area in the image with negative dR/R values falling within 80%, 60%, 40%, or 20% threshold of the strongest pixel intensity. The representative images in Fig. 4 have been cropped with a circular mask to hide skull artifacts such as cement margins, which were excluded from analysis.

### Cytochrome oxidase (CO) staining on flattened tangential sections

Mice were deeply anesthetized with isoflurane and transcardially perfused using 4% paraformaldehyde (PFA) in pH 7.4 0.1M phosphate buffer (PB). The brain was extracted and either left overnight at 4°C in 4% PFA or immediately processed. The hemispheres were separated and subcortical regions were removed to isolate the cortical rind. The cortex was then flattened between two slides using 2-3 mm plastic spacers, and immersed for 48 hours, refrigerated, in 30% sucrose in 0.1 M PB. The flattened cortex was then removed from the slides and sectioned (100 μm) on a freezing microtome. After collection, sections were rinsed in filtered 0.1 M PB 3 times for 5 min each. Sections were then reacted in a 3,3’-diaminobenzidine (DAB) tetrahydrochloride (Sigma D5637-5G) and 0.4 mg/mL cytochrome C (Sigma C-2506) solution over 3-6 hours. Once barrels were visible, the CO reaction was stopped by washing the sections in 0.1M PB. Sections were then mounted and coverslipped.

### Barrel patterning from vGluT2 immunohistochemistry and CO staining

To compare barrel patterning in L4 between genotypes for Fig. 4, flattened sections were prepared, and either stained for CO (as above) or immunostained for VGluT2, which marks thalamocortical axon terminals. VGluT2 staining was performed in tangential flattened cortical sections (50 μm thickness).

Sections were rinsed in 0.1M PB 3 times for 5 min each, blocked for two hours in 10% normal goat serum (NGS) at room temperature, then incubated with primary antibody (guinea pig anti-VGlut2, 1:1000, EMD Millipore Ab2251) overnight at 4°C with 5% NGS and 0.1% Triton X-100. Sections were then washed with PB 3 times for 5 min each, incubated for 2 hours at room temperature with secondary antibody (Alexa Fluor 647 goat anti-guinea pig, 1:500, Invitrogen A21236), and then washed with 1X PB 3 times for 5 min each. Sections were mounted (Vectashield Antifade), counterstained with DAPI (500 nM, Invitrogen D1306) for 12 minutes and coverslipped.

Fluorescence images were acquired using an epifluorescence microscope under a 4x objective. To quantify barrel staining, images from CO or VGluT2-stained sections through L4 were collected using a 4x objective, and tiled to generate a full tangential image across the barrel field. The size of each barrel, and the entire barrel field, were quantified using the polygon measurement tool in FIJI. Sizes for 28 specific barrel columns were measured for each mouse. The total septal area was calculated as total barrel field area - summed area of each individual barrel column. Inter-barrel distance was measured from the center of barrel C1, through the major axis of barrel C2, and ending at the center of barrel C3 in all mice. Only images with clear and distinguishable barrel columns were used in these analyses.

### CRISPRa virus and injections

#### Viral injections

Retro-orbital injections of rAAV-*Scn2a*-CRISPRa reagents were performed as described previously^70^. Viruses were obtained from University of Michigan BRCF Vector Core (RRID:SCR_026696). The titer of each virus was (i) 2.0x10^13^ vg/mL for AAV-PhP.eb-CMV-sadCas9-VP64, (ii) 4.8x10^13^ vg/mL for AAV-PhP.eb-U6-sasgRNA-CMV-mCherry (CRISPRa) vectors, and (iii) 2.0x10^13^ vg/mL titer AAV-PhP.eb-CMV-mCherry (EMPTY). For rAAV-*Scn2a*-CRISPRa injections, 10x10^10^ vg of each virus (i and ii) were injected. For rAAV-EMPTY-CRISPRa injections, 10x10^10^ vg of each virus (i and iii) were injected. For single-vector controls, 10x10^10^ vg of virus (i) were injected, without any virus (ii) or (iii). All injections were suspended in 5% sorbitol (MilliporeSigma S3889), and a total volume of 150 μL was delivered using a 1-mL syringe with 30-gauge needle.

Injections were performed at P30-40 in isoflurane-anesthetized mice warmed on a heating pad. The needle was inserted ∼0.5 inches into the medial canthus underneath the left eyeball until it reached the back of the eye socket. After slowly pushing down the plunger (over 10 seconds), the needle was slowly withdrawn. Mice with signs of pain were administered meloxicam (5–10 mg/kg) for analgesia.

### *In vivo* recordings in CRISPRa-injected mice

Neural recordings and ISOI imaging were performed in CRISPRa injected mice 5-8 weeks after viral injections. A head-post was implanted as in “Headpost Implant Surgery”, above. The C1, C2, and C3 columns in wS1 were localized through the skull using ISOI, and a 0.5-2mm craniotomy was performed. The mouse was transferred to the recording rig, head-fixed and the whiskers inserted into the piezos as described above. A small durotomy was made over the penetration site. Anesthesia was maintained with <0.5% isoflurane and 0.1mg/kg chlorprothixene. A reference ground was attached inside the recording chamber and covered in saline. Recordings in CRISPRa experiments were made in this lightly anesthetized state with Neuropixels 1.0 probes. Probe insertion was as described above. Recording was blind to genotype, and occurred during the dark phase of the light cycle. 3-5 sequential days of recording were performed in each mouse, with the probe located in a different whisker column in S1 on each day. At the end of each day’s recording, the probe was removed, the craniotomy was sealed with silicone sealant (Kwik-Cast, World Precision Instruments), and the recording chamber was sealed with a cover glass (Thomas Scientific 1217N66) and a thin layer of dental cement.

#### Spike data acquisition for Neuropixels recordings

For Neuropixels recordings, spikes were sampled at 30 kHz at 500x gain, common average referenced, and band-pass filtered from 0.3 kHz to 6 kHz. Automated spike sorting was performed with Kilosort3 (https://zenodo.org/records/10713583)^96^, and followed by manual curation using Phy (https://github.com/kwikteam/phy)^98^. Isolated clusters were manually inspected for mean spike waveform amplitude, drift stability, and inter-spike interval refractory period violations, which could not exceed <2% of intervals <1.5 ms. Only well-isolated single units were analyzed. Single units were classified as RS or FS based on trough-to-peak time of the spike waveform at the highest-amplitude recording channel, with a separation criterion of 0.419 ms.

Units in Neuropixels recordings were assigned to layers using current source density (CSD) for each recording penetration, in which the whisker stimulus-evoked local field potential (LFP, 500Hz low-pass) was calculated for each channel. LFP traces were normalized and interpolated between channels (20 um site spacing, 1.6–2x interpolation). CSDs were then calculated using the second spatial derivative and convolved with a 2D (depth × time) Gaussian to isolate whisker-evoked current sources and sinks. The L4-L5A boundary was defined from CSD as the zero-crossing between the most negative current sink (putative L4) and the next deeper current source (putative L5A). Other layers were assigned using lower amplitude current source and sink locations, as well as mouse-averaged anatomical depth estimates relative to the L4/L5a boundary.

#### Analysis of whisker responsiveness and tuning

Analysis of whisker responsiveness and tuning properties for single units in CRISPRa experiments followed the same methods presented above in “Quantitative Analysis of Spike and LFP data”.

#### Localization of recording sites for CRISPRa experiments

Recording penetrations for CRISPRa experiments were localized relative to either VGluT2 staining of barrels, or mCherry expression, which fortuitously revealed the barrel pattern. After the last day of recording, DiI label was introduced at current and prior penetration sites, as localized by surface blood vessels. The brain was removed and fixed, and flattened cortical sections were prepared. Immunohistochemistry was performed for either with fluorophore-tagged streptavidin (streptavidin, Alexa Fluor 647 conjugate, <u>S21374</u>) or mCherry (enhanced by staining with anti-RFP antibody (rat anti-mCherry, 1:800, Invitrogen M11217, and Alexa Fluor goat anti-rat IgG 568, Abcam), using the same methods as described for anti-VGluT2. Di-I marked penetration locations were reconstructed relative to barrel boundaries as described for the awake recording experiments.

### Statistical analysis of summary data

Statistics were performed in MATLAB, Python, or GraphPad Prism. For sample size, N denotes mice, n denotes units or cells. Sample size for each figure panel is given in Table S1. A significance level (alpha) of 0.05 was used throughout. For in vivo recording experiments, statistical results are reported in the Results using an n of units (grouped across mice), but we also verified that all major in vivo effects were consistent across individual mice (Fig. S2 and S7).

For non-normally distributed data, such as in vivo firing rates for populations of units, population medians were compared between groups using permutation tests or the Wilcoxon rank-sum test, and were corrected for multiple comparisons (Bonferroni correction). Permutation tests for differences in median firing rate between two groups are referred to in brief as ‘permutation test’. To test for differences in two population medians with multiple subgroups, we first assessed whether a main effect was present using a Kruskal-Wallis test, and then performed *post hoc* tests (Tukey-Kramer tests), which correct for multiple comparisons. To compare medians from groups of three or more repeated measurements, we used Friedman’s test. One or two-sample Kolmogorov-Smirnov tests for goodness-of-fit were used to test for normality, or to test whether samples came from the same distribution(s).

For normally-distributed data with 2 groups (e.g., slice physiology or barrel topology data), we used unpaired t-tests. For normally-distributed data with >2 groups, we compared population means between groups by first assessing whether a main effect was present using a two-way mixed ANOVA. The chi-squared test of independence was used to compare differences in group counts for categorical data. To test for equivalent medians between two groups (lifetime sparseness for CRISPRa-injected Het and WT in Suppl. Fig. 7), we performed an equivalence test following the procedure in (Lakens, Soc Psych Personality Science 2017) utilizing Mann-Whitney U-tests, using a minimum effect size of 33%.

## ACKNOWLEDGMENTS

This work was supported by R01 MH136475 (K.J.B. and D.E.F.), R01 NS134639 (D.E.F.), and R01 MH125978 (K.J.B.), as well as by grants from the Simons Foundation Autism Research Initiative (SFARI; grant no. 629287: N.A., K.J.B.; grant no. 513133: K.J.B.), the Weill Neurohub Investigator Program (N.A., K.J.B., D.E.F.), and the Medical Research Council Centre of Research Excellence in Therapeutic Genomics (MR/Z504725/1; N.A.). K.V. was supported by a National Science Foundation (NSF) GRFP. N.F.P. was supported by a NSF GRFP and an Autism Science Foundation (ASF) graduate research fellowship.

## Supplemental Figures

**Figure S1.**
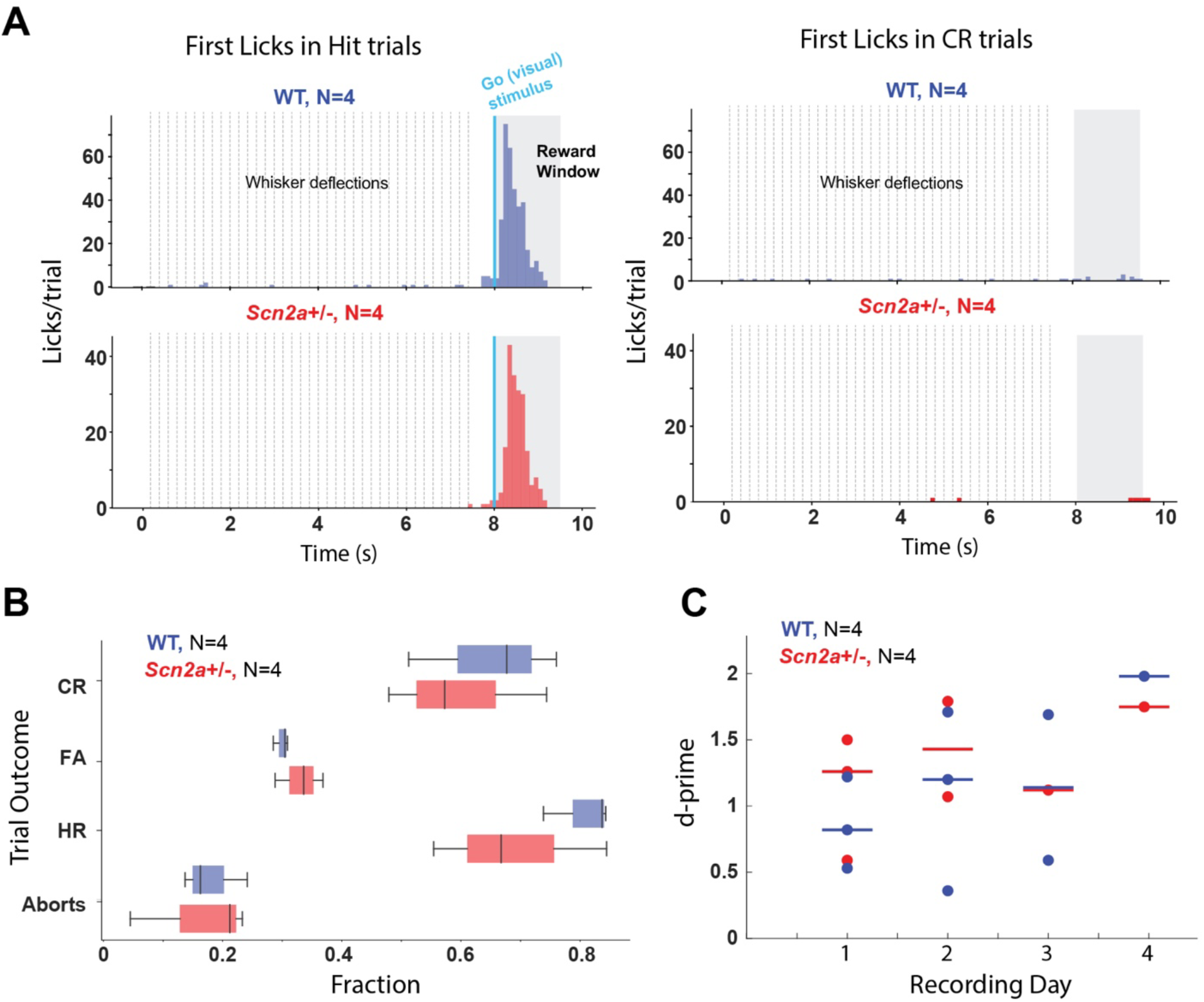
Task engagement in wild type and *Scn2a^+/−^* mice. **A)** Histogram of first licks in the behavioral task, relative to whisker stimulation and reward window onset. Left: Hit trials (licks in reward window after blue light flash), right: correct rejections (CR, no licks in reward window above threshold, with no blue light flash). Dashed lines: whisker deflections. N=4 WT, N=4 Scn2aHet mice. **B**) Average proportion of correct rejections (CRs), false alarms (FAs), hits (HR), and aborted trials (aborts) across mice. WT: blue, Scn2aHet: red. Box: inter-quartile range. Dot=mouse. **C**) Behavioral performance during visual detection task across recording days, per mouse. Dashed line: chance level. Solid line: median, dot: mouse.

**Figure S2:**
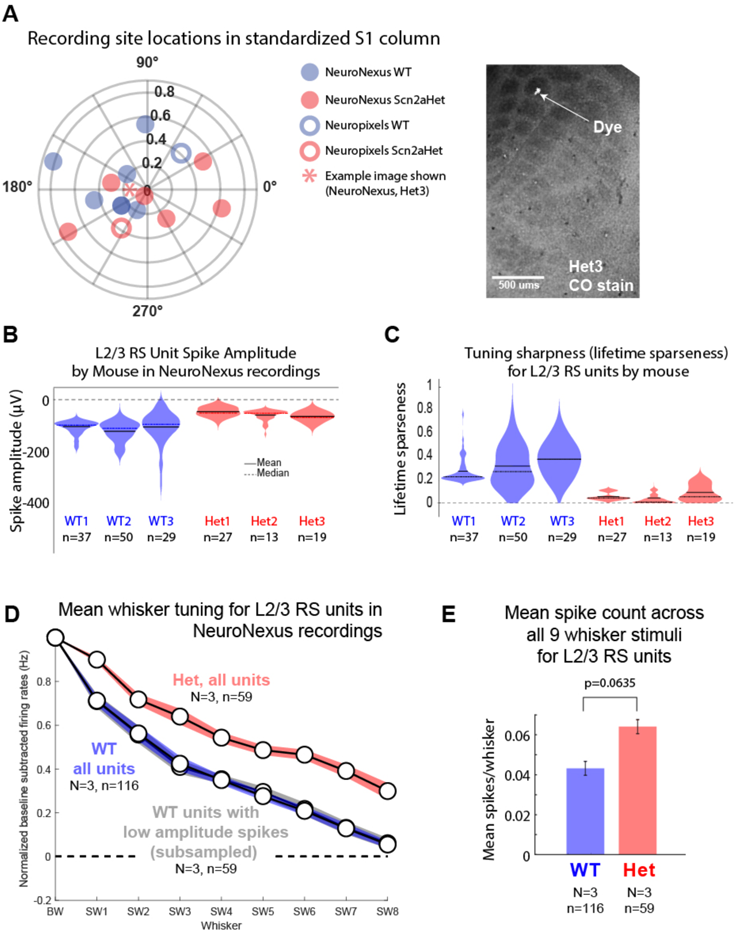
Recording locations and additional analysis of L2/3 RS spike waveforms and tuning properties from NeuroNexus probe recordings in awake mice. **A**) Left: Penetration sites for each recording site (from NeuroNexus and Neuropixels recordings) within a standardized barrel column. Radius, penetration distance between barrel column center relative to length of the major column axis. Right: Example of penetration (dye mark) in the B1 column (corresponding to star in polar plot). **B**) Distribution of spike amplitude for L2/3 RS units in each mouse. n=units. **C)** Lifetime sparseness for all L2/3 RS units in Fig. 2F, by mouse. Each *Scn2a+/−* mouse showed broad whisker tuning (low lifetime sparseness). **D)** Mean rank-ordered whisker tuning across whisker-responsive L2/3 RS units, for all units in WT mice, all units in Het mice, or a subsample of WT units that are matched for extracellular spike amplitude to the Het units. Each unit’s tuning curve was normalized to that unit’s BW response, then baseline- subtracted, and then averaged across units. Dashed line=spontaneous spike rate. **E)** Mean whisker-evoked spiking for L2/3 RS units in NeuroNexus recordings, calculated across all 9 whiskers. Same units as in Fig. 2C.

**Figure S3:**
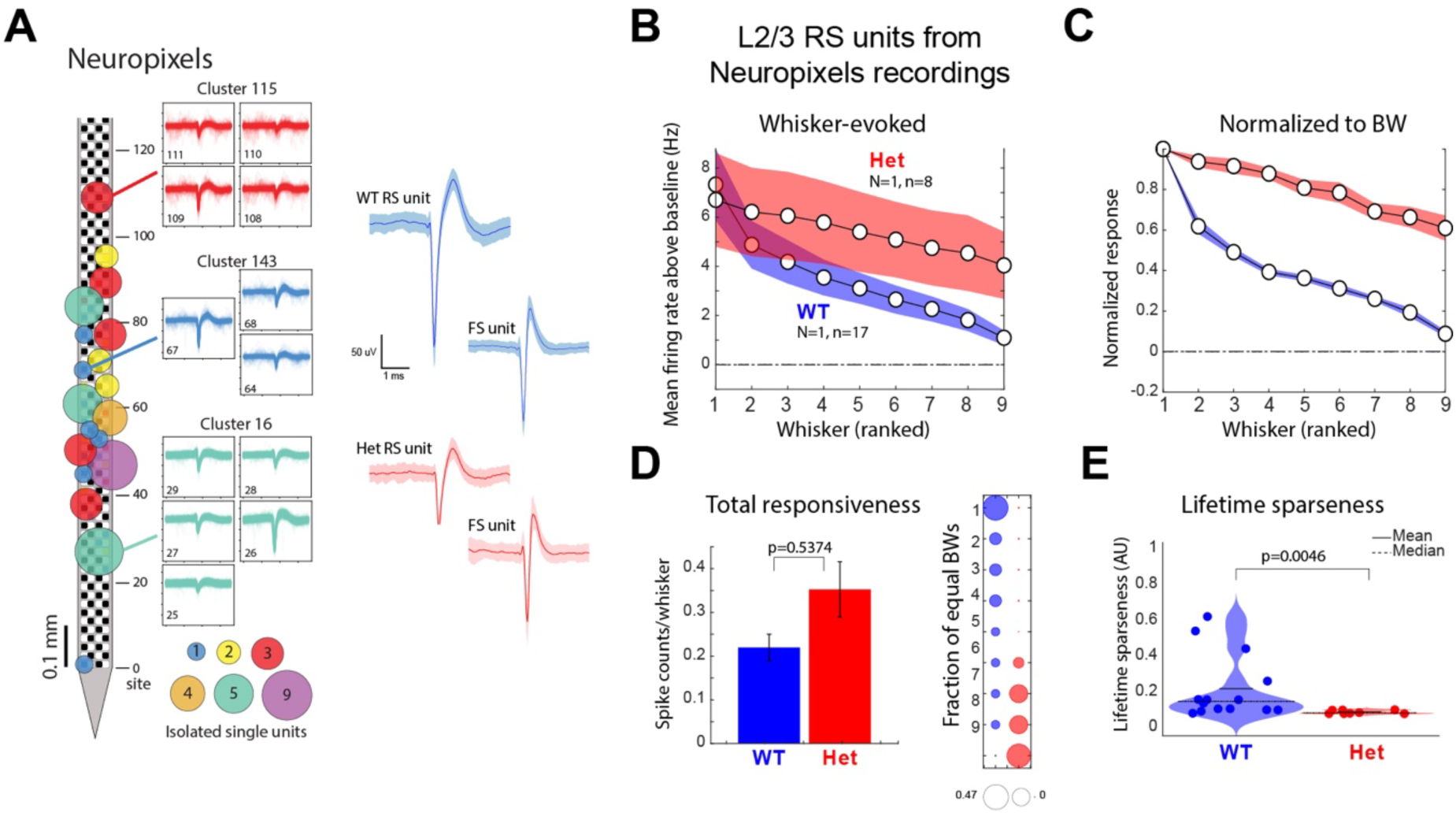
L2/3 RS unit properties from Neuropixels recordings. **A**) Left: Neuropixels probe with example isolated RS and FS units from WT (blue) and Scn2aHet (red) mice. Line=mean, shaded region=SEM. **B**) Mean rank-ordered, baseline-subtracted whisker tuning curve for L2/3 RS units in 1 WT mouse and 1 Scn2a+/− mouse recorded with Neuropixels. Dashed line: spontaneous firing. All error shading/lines are 68% CI. **C**) Same data as (B), normalized to the BW response of each unit before averaging. Conventions as in Fig. 2D. **D**) Left: Average whisker-evoked firing rate across all 9 whisker stimuli, for each L2/3 RS unit in the 2 genotypes. Wilcoxon rank-sum test. Right: number of equivalent BWs in L2/3 RS units. Circle size indicates fraction of units. WT: blue, Scn2aHet: red. Chi-squared, p=4.16e-05, alpha=0.05. **E**) Quantification of tuning sharpness by lifetime sparseness for L2/3 RS units in these 2 mice. Each dot is one unit. Solid line: mean, dashed line: median. Wilcoxon rank-sum test. Alpha was 0.05 for all tests.

**Figure S4:**
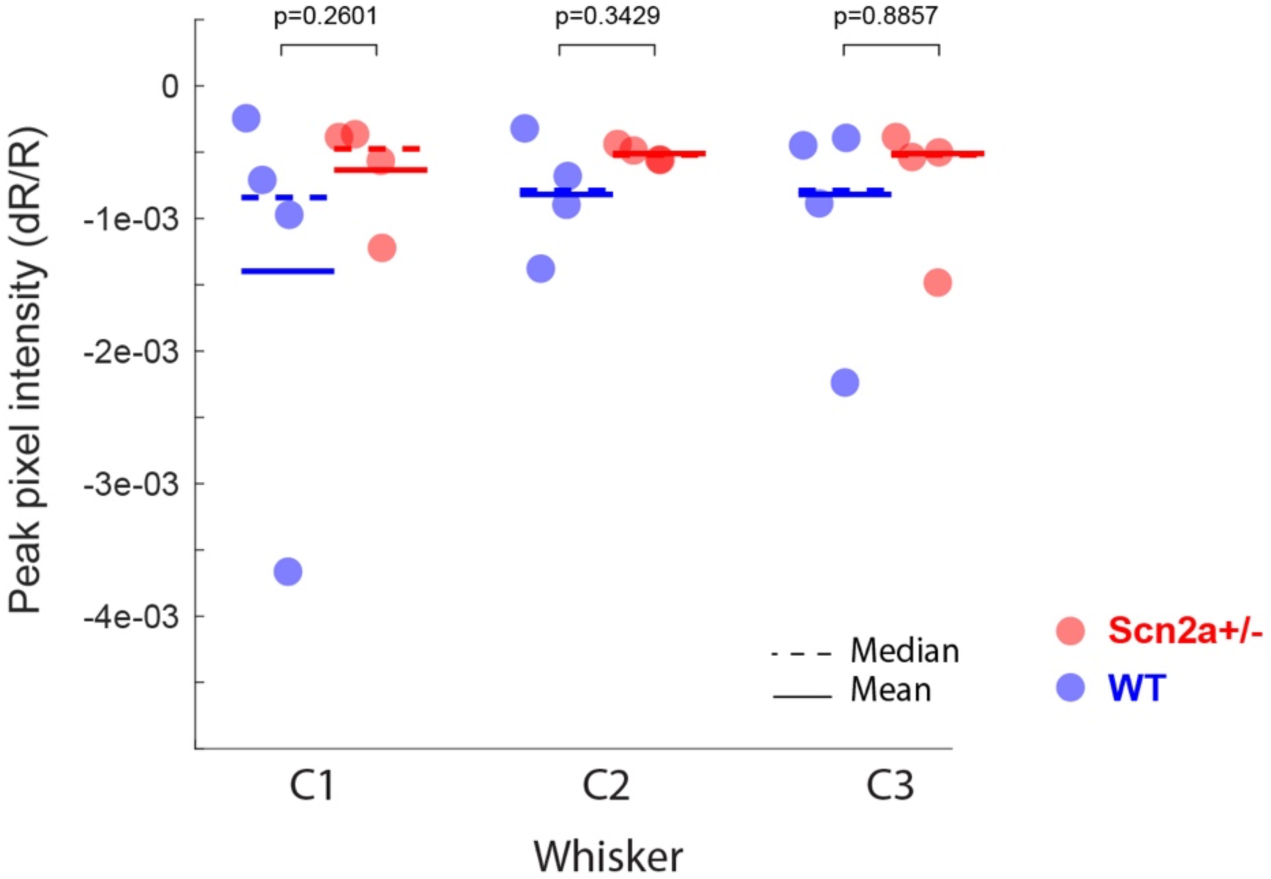
Peak pixel intensity in ISOI does not differ between genotypes. Greatest pixel intensity measured during ISOI per mouse, by whisker. WT=blue, Scn2aHet=red. WT: N=4 mice, n=4 ROIs per whisker. Scn2aHet: N=4 mice, n=4 ROIs per whisker. Median dR/R for each condition was: WT C1: −0.0007 ± 0.0014, C2: −0.0008 ± 0.0007, C3: −0.0007 ± 0.0013; *Scn2a^+/−^* C1: −0.0005 ± 0.0008, C2: −0.0005 ± 0.00009, C3: −0.0005 ± 0.0006. The median dR/R did not differ between genotypes for any whisker. Statistics: Wilcoxon rank-sum tests, alpha=0.0167 after Bonferroni correction for multiple comparisons.

**Figure S5:**
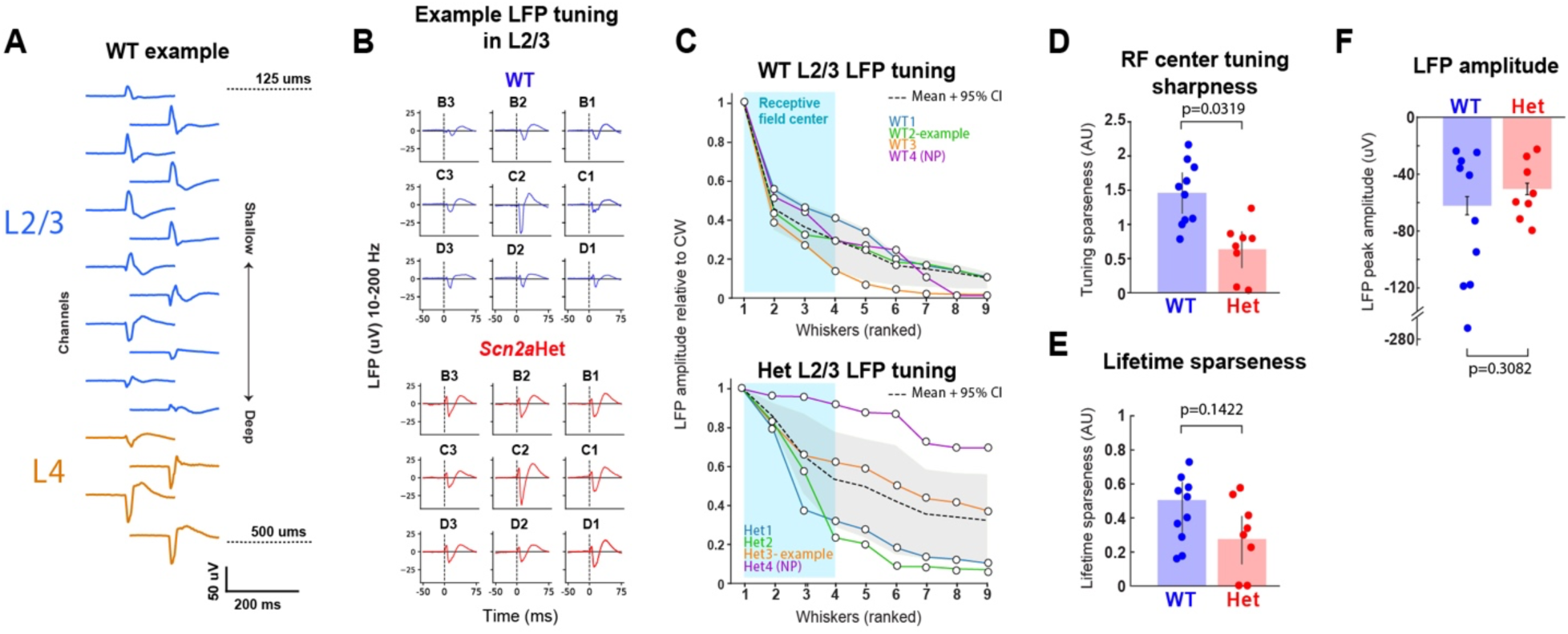
Somatotopic blurring in local field potential (LFP) recordings in *Scn2a^+/−^* mice. **A)** Example LFP traces evoked by CW deflection on a 16-channel probe during the behavioral task. Orange: L4 channels, blue: L2/3 channels. Laminar labels were determined based on the geometric mean depth of each 4-channel group, relative to channel depth below the pia (dotted lines). **B**) LFP waveforms (10-200 Hz bandpass) from L2/3 in one WT and one Scn2aHet mouse, in response to 9 single-whisker stimuli. Both penetrations were in the C2 column. WT example shown is from channel 9 (ninth down from top) in A. **C**) Rank ordered LFP tuning curve for 4 WT and 4 Scn2aHet mice, based on LFP amplitude normalized to the CW response. Each line is mean across all recording sites in one mouse. Dashed line and grey region are mean and 95% CI. Example refers to data shown in B. Recordings were made with NeuroNexus probes, except for one mouse in each group recorded with NeuroPixels (NP). Blue shaded region indicates whisker receptive field center. WT: N=4 mice, n=10 sites. Scn2aHet: N=4 mice, 8 sites. Permutation tests and alpha=0.05 used for all tests. **D**) Receptive field (RF) center tuning sharpness in L2/3 LFP. Each dot is one penetration site and error bar is 95% CI. **E**) Lifetime sparseness per penetration site. **F**) LFP amplitude for the strongest whisker, for each penetration site.

**Figure S6:**
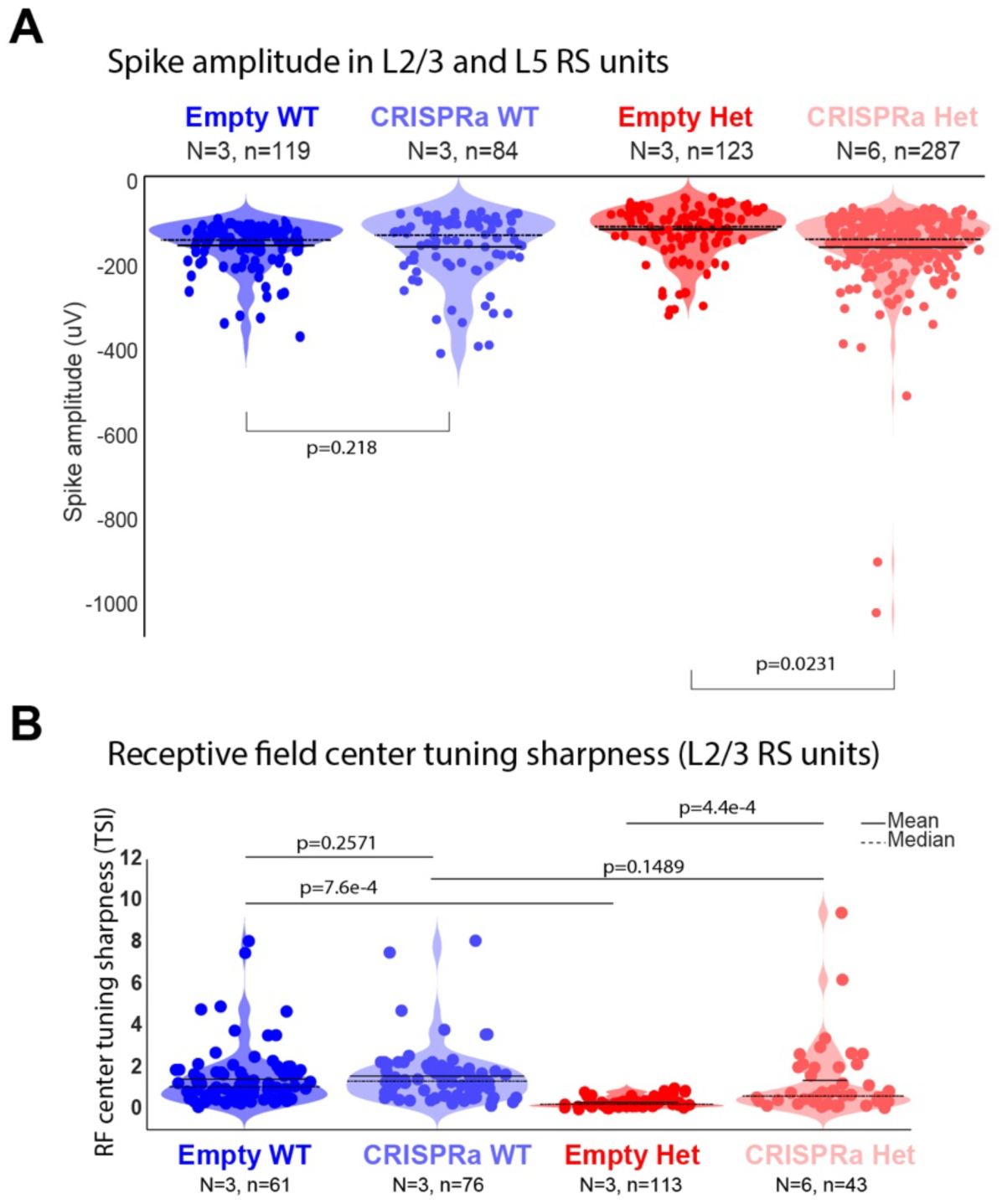
Additional analysis of spike properties and tuning in mice in the CRISPRa experiment. **A**) Spike amplitude for L2/3 C L5 RS units for four experimental groups. Dashed line=median, solid line=mean. Wilcoxon rank-sum tests, alpha=0.025 for multiple comparisons. All sample sizes are N=mice, n=single units. **B**) RF center tuning sharpness index (TSI) for L2/3 RS units. Tukey-Kramer tests, alpha=0.05.

**Figure S7:**
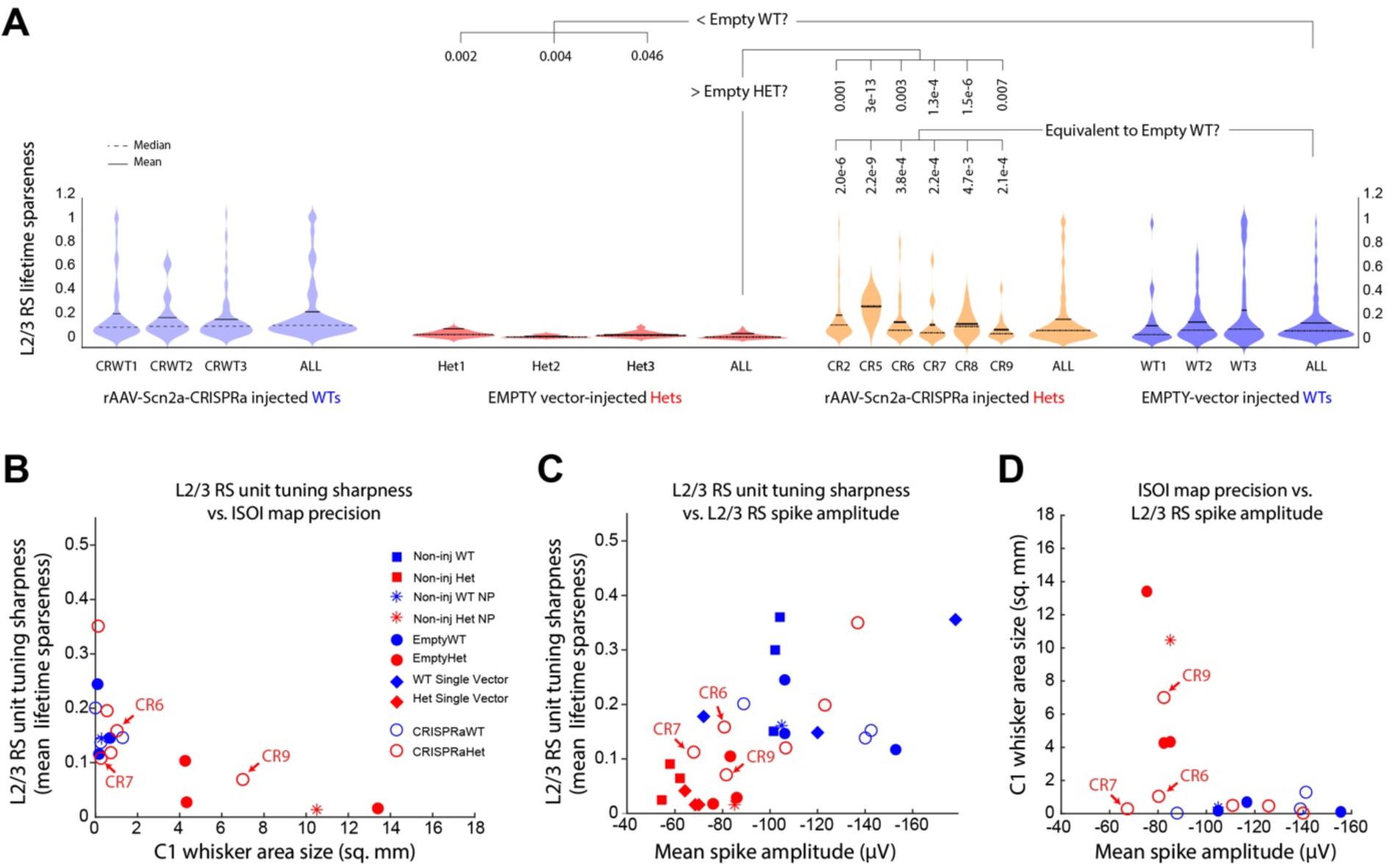
Relationship of L2/3 RS somatotopic tuning, ISOI map topography, and L2/3 RS spike amplitude in each mouse. **A**) Lifetime sparseness for all L2/3 RS units in each mouse, per experimental group in the CRISPRa experiment. Each violin plot is an individual mouse. Statistics are from permutation tests between each indicated mouse and all empty-vector injected WTs, with FDR correction for multiple comparisons. Alpha=0.05. To test for equivalent medians between two groups (lifetime sparseness for CRISPRa-injected Hets and WT in Suppl. Fig. 7), we performed an equivalence test following the procedure in (Lakens, Soc Psych Personality Science 2017) utilizing Mann-Whitney U-tests, using a minimum effect size of 33%. Within each family of comparisons, reported p-values have been adjusted for multiple comparisons by Storey’s method (Storey, 2022). **B**) Relationship of L2/3 RS tuning sharpness (lifetime sparseness) from each mouse with C1 whisker area size. Filled symbols and asterisks are mice without rAAV-CRISPRa-Scn2a (either uninjected, Empty vector injected, or single vector control mice). Open symbols are mice with rAAV-CRISPRa-Scn2a injections. In general, mice with sharp ISOI maps (low area) had sharpest single-unit tuning in L2/3 RS units. **C**) Mean L2/3 RS unit lifetime sparseness by mean L2/3 RS unit spike amplitude per mouse. **D**) Mean C1 whisker area size by mean L2/3 RS spike amplitude per mouse. In B-D, each point is one mouse.

**Figure S8.**
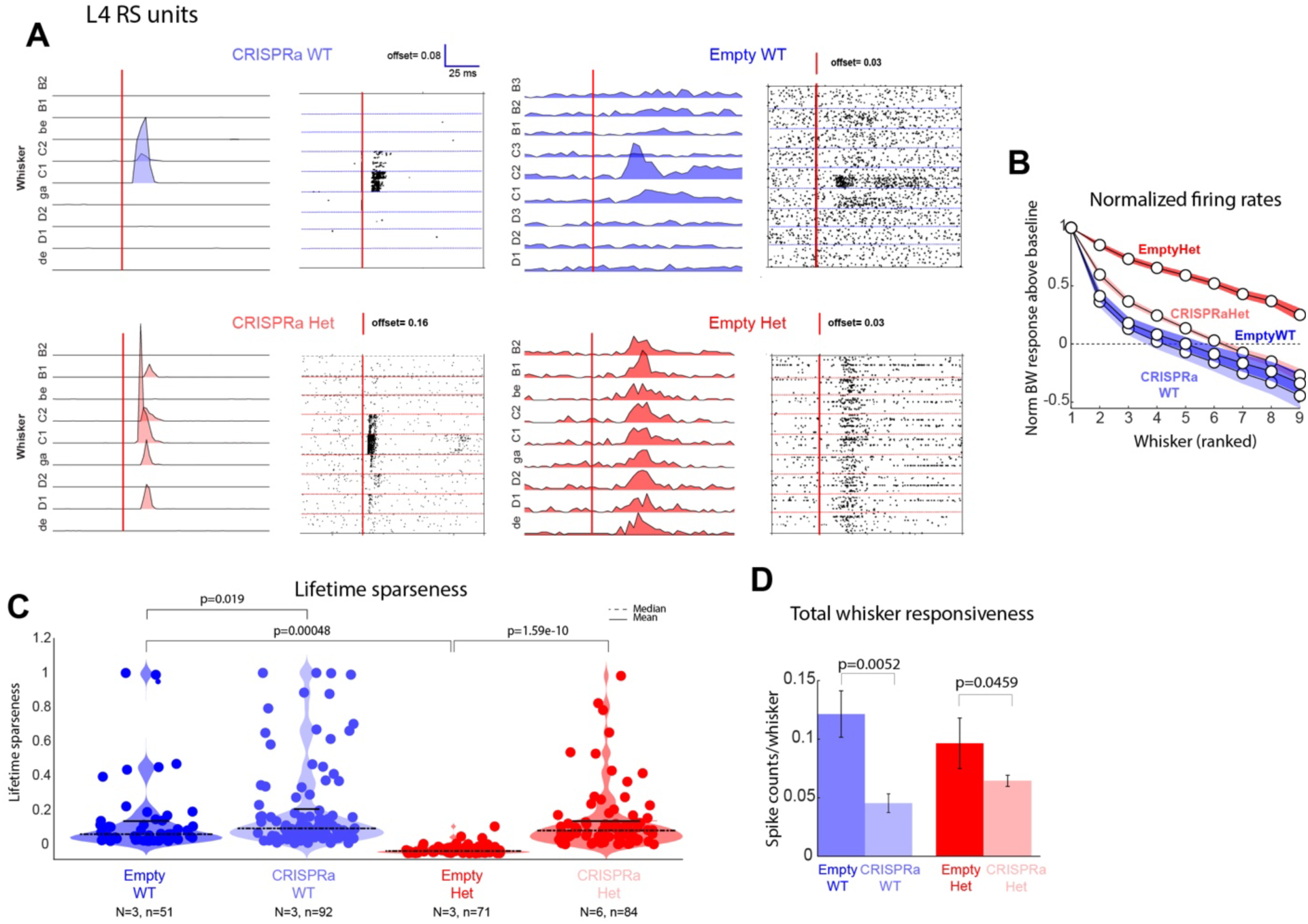
CRISPRa rescues degraded somatotopic tuning for L4 RS units. **A**) 9 whisker somatotopic receptive fields in L4 RS units. Vertical line: whisker deflection. CRISPRa: mice injected with full dual-vector strategy in A. Empty: mice injected with one vector lacking the Scn2a guide RNA in A. **B**) Mean rank-ordered and baseline-subtracted whisker tuning curves. Dashed line: spontaneous activity. All error bars are 68% CI. **C**) Lifetime sparseness. Statistics are from Tukey-Kramer test. Alpha=0.05. N=mice, n=units. **D**) Total sensory responsiveness (cumulative whisker-evoked spiking/nine whisker stimuli). Statistics are from Wilcoxon rank-sum tests and alpha=0.025 for multiple comparisons.

**Figure S9.**
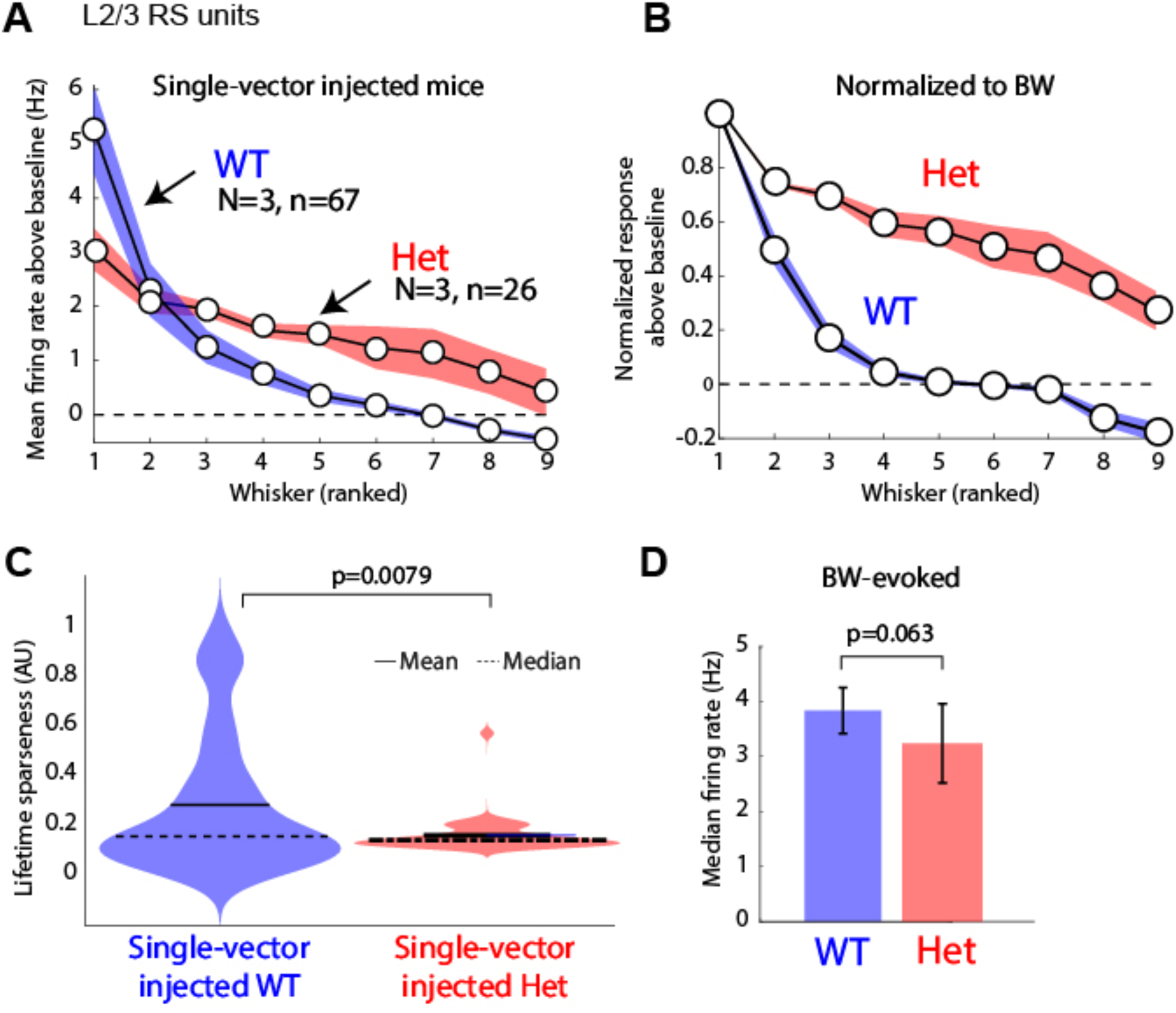
Sensory tuning in L2/3 RS units from mice in single-vector control experiments. **A**) Mean rank-ordered, baseline-subtracted whisker tuning curves from L2/3 RS units from mice injected with only one viral vector (AAV-PhP.eb-CMV-SadCas9-VP64). These mice (N=3 WT, N=3 Scn2a+/−) did not receive the guide RNA vector, and thus upgregulation of *Scn2a* will be absent. Dashed line: spontaneous firing rate. All error bars are 68% CI. **B**)) Same as in A, but with tuning curves normalized to each unit’s BW response before averaging. **C**) Lifetime sparseness for the units in A-B. Statistics from Tukey-Kramer test. Alpha=0.05. N=mice, n=units. **D**) Bootstrapped median firing rate for the same units as in A-C. Error bars are 68% CI. Statistics are from permutation test.

**Figure S10.**
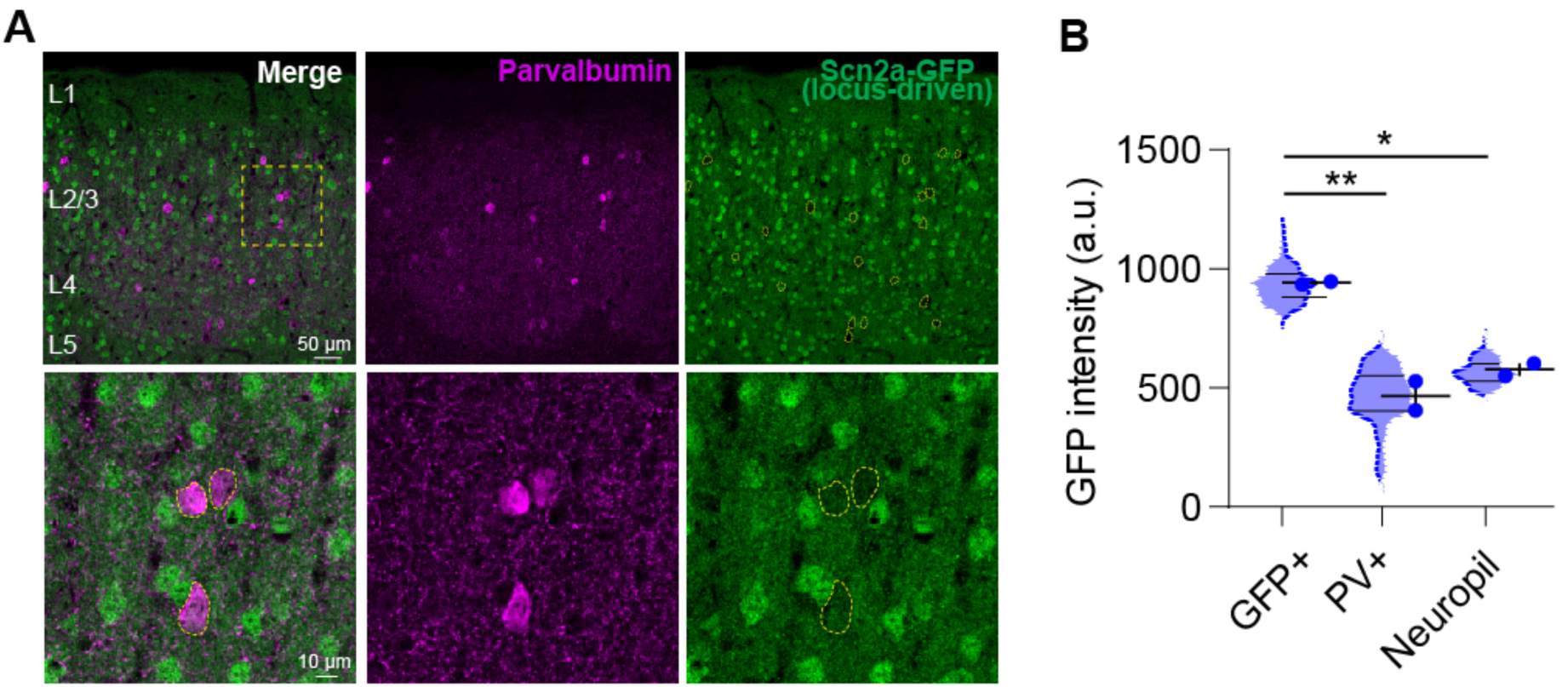
PV cells lack expression of GFP from the endogenous Scn2a locus in Scn2a+/KI mice. **A)** S1 section from a Scn2a+/KI mouse (Tamura et al., Nature 2025) that expresses GFP from the endogenous Scn2a locus, and was immunostained for PV. Green, GFP fluorescence. Bottom row, higher magnification from the small box on top. PV interneurons (dashed outlines) show no GFP fluorescence. **B)** Quantification of mean GFP intensity in each cell. GFP+ indicates GFP+, PV- somata. PV+ indicates PV+ somata. Violin plots show intensity distribution across all cells in the category. Circles are cell means per mouse (N=2 mice). Horizontal bar, mean across mice. Statistics from nested one-way ANOVA, F(2,3) = 41.61, **p = 0.0065, n = 2 mice, 50-80 ROIs per group. Post-hoc tests show that GFP intensity is greater in non-PV cells than PV cells, and that PV cells are not different than neuropil.

**Table S1.** Sample size for each measurement.

| **Fig. Panel** | **Experimental Group** | **N, n (mice, cells) or other units listed below** | **Group composition** |
| --- | --- | --- | --- |
| 1B-C | WT<br>Het | 3, 8<br>3, 9 | L2/3 PYR cells |
| 1D | WT<br>Het | 4, 13<br>4, 11 | L2/3 PYR cells |
| 1F | WT<br>Het | 3, 327<br>3, 161 | All L2/3 RS and FS units |
| 1G, L | WT<br>Het | 3, 259<br>3, 109 | All L2/3 RS units |
| 1H-K | WT<br>Het | 3, 116<br>3, 59 | Whisker-responsive L2/3 RS units |
| 1M-O | WT<br>Het | 3, 106<br>3, 32 | Whisker-responsive L2/3 FS units |
| 1P | WT<br>Het | 3, 111<br>3, 49 | All L2/3 FS units |
| 2B | WT<br>Het | 3, 77<br>3, 59 | L2/3 RS units that were tuned for the CW |
| 2C-F | WT<br>Het | 3, 116<br>3, 59 | Whisker-responsive L2/3 RS units (same as 1H-K) |
| 3C | WT<br>Het | 4 mice, 4 WRAs per panel<br>4 mice, 4 WRAs per panel | All whisker response areas across mice |
| 4A-D | WT<br>Het | 3, 74<br>3, 33 | Whisker-responsive L4 RS units |
| 4E-I | WT<br>Het | 3, 74<br>3, 33 | L4 RS units that were tuned for the CW |
| 4L | WT (CO)<br>Het (CO)<br>WT (VGluT2)<br>Het (VGluT2) | 4 mice, 76 barrels<br>4 mice, 76 barrels<br>3 mice, 84 barrels<br>3 mice, 84 barrels | CO-stained barrels<br>CO-stained barrels<br>VGluT2-stained barrels<br>VGluT2-stained barrels |
| 4M | WT (CO)<br>Het (CO)<br>WT (VGluT2)<br>Het (VGluT2) | 4 mice, 4 fields<br>4 mice, 4 fields<br>3 mice, 3 fields<br>3 mice, 3 fields | CO-stained barrel fields<br>CO-stained barrel fields<br>VGluT2-stained barrel fields<br>VGluT2-stained barrel fields |
| 5A | WT<br>Het | 3, 116<br>3, 59 | Whisker-responsive L2/3 RS units (same as 1H-K) |
| 5C-G | WT<br>Het | 3, 59<br>3, 59 | L2/3 RS units that were tuned for the CW,<br>subsampled for equal population size in WT and<br>HET |
| 6C-H | EmptyWT<br>CRISPRaWT<br>EmptyHet<br>CRISPRaHet | 3, 61<br>3, 76<br>3, 72<br>6, 43 | Whisker-responsive L2/3 RS units |
| 7B | Non-inj WT<br>EmptyWT<br>CRISPRaWT<br>Non-inj Het<br>EmptyHet<br>CRISPRaHet | 4 mice, 4 WRAs<br>3 mice, 3 WRAs<br>3 mice, 3 WRAs<br>4 mice, 4 WRAs<br>3 mice, 3 WRAs<br>6 mice, 6 WRAs<br>(per panel) | Whisker activation areas (data for non-injected<br>mice are the same as shown in 3C) |
| 8B | WT (PV+)<br>WT (Slc17a7+)<br>Het (PV+)<br>Het (Slc17a7+) | 3 mice, 10 cells<br>3 mice, 12 cells<br>3 mice, 11 cells<br>3 mice, 11 cells |  |
| 8E-F | WT<br>Het | 3 mice, 9 cells<br>3 mice, 9 cells | S5E2+ fast-spiking cells in L2/3 of S1 |
| 8H-J | WT<br>Het | 6 mice, 17 cells<br>5 mice, 16 cells | S5E2+ fast-spiking cells<br>in L2/3 of S1 |
| 8K | EmptyWT<br>CRISPRaWT<br>EmptyHet<br>CRISPRaHet | 3, 38<br>3, 19<br>3, 41<br>6, 93 | Whisker-responsive L2/3 and L4 FS units |
| S1A-C | WT<br>Het | 4 mice, 10 sessions<br>4 mice, 8 sessions | Behavioral sessions |
| S2A | WT<br>Het | 4 mice, 10<br>penetrations<br>4 mice, 8<br>penetrations | NeuroNexus and Neuropixels recording<br>penetrations |
| S2B-C | individual mice:<br>WT1, 2, 3<br>Het1, 2, 3 | 37, 50, and 29 units<br>27, 13, and 19 units | Whisker-responsive L2/3 RS units (same as 1H-K) |
| S2D | WT<br>Het<br>WT subsampled | 3, 116<br>3, 59<br>3, 59 | Whisker-responsive L2/3 RS units (same as 1H-K) |
| S2E | WT<br>Het | 3, 116<br>3, 59 | Whisker-responsive L2/3 RS units (same as 1H-K) |
| S3B-E | WT<br>Het | 1, 17<br>1, 8 | Whisker-responsive L2/3 RS units in Neuropixels<br>recordings |
| S4 | WT<br>Het | 4 mice, 4 WRAs<br>4 mice, 4 WRAs | Peak ISOI pixel intensity for each WRA in each<br>mouse |
| S5C | WT<br>Het | 4 mice<br>4 mice | Each curve is mean LFP tuning recorded in each individual mouse |
| S5C-F | WT<br>Het | 4 mice, 10 recordings<br>4 mice, 8 recordings | All L2/3 LFP recordings from NeuroNexus and Neuropixels experiments |
| S6A | EmptyWT<br>CRISPRaWT<br>EmptyHet<br>CRISPRaHet | 3, 119<br>3, 84<br>3, 123<br>6, 287 | All L2/3 and L5 RS units |
| S6B | EmptyWT<br>CRISPRaWT<br>EmptyHet<br>CRISPRaHet | 3, 61<br>3, 76<br>3, 113<br>6, 43 | Whisker-responsive L2/3 RS units |
| S7A | individual mice:<br>CRWT1, 2, 3<br>Het1, 2, 3<br>CR2, 5, 6, 7, 8, 9<br>WT1, 2, 3 | 43, 19, 14 units<br>40, 34, 45 units<br>9, 6, 10, 5, 7, 6 units<br>11, 28, 22 units | Whisker-responsive L2/3 RS units |
| S7B-D |  | Each point is an individual mouse | Mean tuning of whisker-responsive units, mean area of WRA |
| S8B-D | EmptyWT<br>CRISPRaWT<br>EmptyHet<br>CRISPRaHet | 3, 51<br>3, 92<br>3, 71<br>6, 84 | Whisker-responsive L4 RS units |
| S9A-E | SingleWT<br>SingleHet | 3, 67<br>3, 26 | Whisker-responsive L2/3 RS units in single vector-injected control mice |

**Table S2:** Whisker-evoked ISOI magnitude (dR/R) at peak pixel. Values are whisker-evoked dR/R (mean ± SEM across mice) relative to baseline, at the peak pixel (darkest pixel).

|  | C1 | C2 | C3 | Mouse N |
| --- | --- | --- | --- | --- |
| EmptyWT | 0.9997 ± 0.0003 | 0.9996 ± 0.0002 | 0.9996 ± 0.00008 |  |
| EmptyHet | 0.9997 ± 0.0002 | 0.9997 ± 0.00005 | 0.9997 ± 0.0003 |  |
| CRISPRaWT | 0.9997 ± 0.0002 | 0.9998 ± 0.0002 | 0.9996 ± 0.0002 |  |
| CRISPRaHet | 0.9997 ± 0.0001 | 0.9997 ± 0.00005 | 0.9997 ± 0.0003 |  |

## REFERENCES

1. Ben-Shalom, R., Keeshen, C.M., Berrios, K.N., An, J.Y., Sanders, S.J., and Bender, K.J. (2017). Opposing Effects on NaV1.2 Function Underlie Differences Between SCN2A Variants Observed in Individuals With Autism Spectrum Disorder or Infantile Seizures. Biol Psychiatry 82, 224–232. 10.1016/j.biopsych.2017.01.009.

2. Begemann, A., Acuña, M.A., Zweier, M., Vincent, M., Steindl, K., Bachmann-Gagescu, R., Hackenberg, A., Abela, L., Plecko, B., Kroell-Seger, J., et al. (2019). Further corroboration of distinct functional features in SCN2A variants causing intellectual disability or epileptic phenotypes. Mol Med 25, 6. 10.1186/s10020-019-0073-6.

3. Wolff, M., Johannesen, K.M., Hedrich, U.B.S., Masnada, S., Rubboli, G., Gardella, E., Lesca, G., Ville, D., Milh, M., Villard, L., et al. (2017). Genetic and phenotypic heterogeneity suggest therapeutic implications in SCN2A-related disorders. Brain 140, 1316–1336. 10.1093/brain/awx054.

4. Sanders, S.J., Campbell, A.J., Cottrell, J.R., Moller, R.S., Wagner, F.F., Auldridge, A.L., Bernier, R.A., Catterall, W.A., Chung, W.K., Empfield, J.R., et al. (2018). Progress in Understanding and Treating SCN2A- Mediated Disorders. Trends Neurosci 41, 442–456. 10.1016/j.tins.2018.03.011.

5. Thomas, B.R., Ludwig, N.N., Pelletier, D., Bauer, M., Hommer, R., Smith-Hicks, C., and O’Connor, J.T. (2024). Cortical Vision Impairment (CVI)-informed assessment and treatment of challenging behavior in a child with SCN2A-related disorder. J Neurodev Disord 16, 66. 10.1186/s11689-024-09580-7.

6. Berg, A.T., Thompson, C.H., Myers, L.S., Anderson, E., Evans, L., Kaiser, A.J.E., Paltell, K., Nili, A.N., DeKeyser, J.-M.L., Abramova, T.V., et al. (2024). Expanded clinical phenotype spectrum correlates with variant function in SCN2A-related disorders. Brain 147, 2761–2774. 10.1093/brain/awae125.

7. Sanders, S.J., He, X., Willsey, A.J., Ercan-Sencicek, A.G., Samocha, K.E., Cicek, A.E., Murtha, M.T., Bal, V.H., Bishop, S.L., Dong, S., et al. (2015). Insights into Autism Spectrum Disorder Genomic Architecture and Biology from 71 Risk Loci. Neuron 87, 1215–1233. 10.1016/j.neuron.2015.09.016.

8. Satterstrom, F.K., Kosmicki, J.A., Wang, J., Breen, M.S., De Rubeis, S., An, J.-Y., Peng, M., Collins, R., Grove, J., Klei, L., et al. (2020). Large-Scale Exome Sequencing Study Implicates Both Developmental and Functional Changes in the Neurobiology of Autism. Cell 180, 568–584.e23. 10.1016/j.cell.2019.12.036.

9. Wang, T., Kim, C.N., Bakken, T.E., Gillentine, M.A., Henning, B., Mao, Y., Gilissen, C., The SPARK Consortium, Nowakowski, T.J., and Eichler, E.E. (2022). Integrated gene analyses of de novo variants from 46,612 trios with autism and developmental disorders. Proceedings of the National Academy of Sciences 119, e2203491119. 10.1073/pnas.2203491119.

10. Willsey, H.R., Willsey, A.J., Wang, B., and State, M.W. (2022). Genomics, convergent neuroscience and progress in understanding autism spectrum disorder. Nat Rev Neurosci 23, 323–341. 10.1038/s41583-022-00576-7.

11. Hudac, C.M., Friedman, N.R., Ward, V.R., Estreicher, R.E., Dorsey, G.C., Bernier, R.A., Kurtz-Nelson, E.C., Earl, R.K., Eichler, E.E., and Neuhaus, E. (2024). Characterizing Sensory Phenotypes of Subgroups with a Known Genetic Etiology Pertaining to Diagnoses of Autism Spectrum Disorder and Intellectual Disability. J Autism Dev Disord 54, 2386–2401. 10.1007/s10803-023-05897-9.

12. Schust, L.F., Burke, J., SanInocencio, C., Bryan, B.A., Ho, K.S., and Egan, S.M. (2024). A patient organization perspective: charting the course to a cure for SCN2A-related disorders. Ther Adv Rare Dis 5, 26330040241292645. 10.1177/26330040241292645.

13. Balasco, L., Provenzano, G., and Bozzi, Y. (2020). Sensory Abnormalities in Autism Spectrum Disorders: A Focus on the Tactile Domain, From Genetic Mouse Models to the Clinic. Front Psychiatry 10, 1016. 10.3389/fpsyt.2019.01016.

14. Baranek, G.T., Watson, L.R., Boyd, B.A., Poe, M.D., David, F.J., and McGuire, L. (2013). Hyporesponsiveness to social and nonsocial sensory stimuli in children with autism, children with developmental delays, and typically developing children. Dev Psychopathol 25, 307–320. 10.1017/S0954579412001071.

15. Coskun, M.A., Varghese, L., Reddoch, S., Castillo, E.M., Pearson, D.A., Loveland, K.A., Papanicolaou, A.C., and Sheth, B.R. (2009). How somatic cortical maps differ in autistic and typical brains. Neuroreport 20, 175–179. 10.1097/WNR.0b013e32831f47d1.

16. Hu, W., Tian, C., Li, T., Yang, M., Hou, H., and Shu, Y. (2009). Distinct contributions of Na(v)1.6 and Na(v)1.2 in action potential initiation and backpropagation. Nat Neurosci 12, 996–1002. 10.1038/nn.2359.

17. Li, T., Tian, C., Scalmani, P., Frassoni, C., Mantegazza, M., Wang, Y., Yang, M., Wu, S., and Shu, Y. (2014). Action Potential Initiation in Neocortical Inhibitory Interneurons. PLOS Biology 12, e1001944. 10.1371/journal.pbio.1001944.

18. Spratt, P.W.E., Alexander, R.P.D., Ben-Shalom, R., Sahagun, A., Kyoung, H., Keeshen, C.M., Sanders, S.J., and Bender, K.J. (2021). Paradoxical hyperexcitability from NaV1.2 sodium channel loss in neocortical pyramidal cells. Cell Rep 36, 109483. 10.1016/j.celrep.2021.109483.

19. Spratt, P.W.E., Ben-Shalom, R., Keeshen, C.M., Burke, K.J., Clarkson, R.L., Sanders, S.J., and Bender, K.J. (2019). The Autism-Associated Gene Scn2a Contributes to Dendritic Excitability and Synaptic Function in the Prefrontal Cortex. Neuron 103, 673–685.e5. 10.1016/j.neuron.2019.05.037.

20. Shin, W., Kweon, H., Kang, R., Kim, D., Kim, K., Kang, M., Kim, S.Y., Hwang, S.N., Kim, J.Y., Yang, E., et al. (2019). Scn2a Haploinsufficiency in Mice Suppresses Hippocampal Neuronal Excitability, Excitatory Synaptic Drive, and Long-Term Potentiation, and Spatial Learning and Memory. Front Mol Neurosci 12, 145. 10.3389/fnmol.2019.00145.

21. Robertson, C.E., and Baron-Cohen, S. (2017). Sensory perception in autism. Nat Rev Neurosci 18, 671– 684. 10.1038/nrn.2017.112.

22. Schaffler, M.D., Middleton, L.J., and Abdus-Saboor, I. (2019). Mechanisms of Tactile Sensory Phenotypes in Autism: Current Understanding and Future Directions for Research. Curr Psychiatry Rep 21, 134. 10.1007/s11920-019-1122-0.

23. Rubenstein, J.L.R., and Merzenich, M.M. (2003). Model of autism: increased ratio of excitation/inhibition in key neural systems. Genes Brain Behav 2, 255–267. 10.1034/j.1601-183x.2003.00037.x.

24. Markram, K., and Markram, H. (2010). The intense world theory - a unifying theory of the neurobiology of autism. Front Hum Neurosci 4, 224. 10.3389/fnhum.2010.00224.

25. Dinstein, I., Heeger, D.J., Lorenzi, L., Minshew, N.J., Malach, R., and Behrmann, M. (2012). Unreliable evoked responses in autism. Neuron 75, 981–991. 10.1016/j.neuron.2012.07.026.

26. Bhaskaran, A.A., Gauvrit, T., Vyas, Y., Bony, G., Ginger, M., and Frick, A. (2023). Endogenous noise of neocortical neurons correlates with atypical sensory response variability in the Fmr1(-/y) mouse model of autism. Nat Commun 14, 7905. 10.1038/s41467-023-43777-z.

27. Haigh, S.M. (2018). Variable sensory perception in autism. Eur J Neurosci 47, 602–609. 10.1111/ejn.13601.

28. Zhang, Y., Bonnan, A., Bony, G., Ferezou, I., Pietropaolo, S., Ginger, M., Sans, N., Rossier, J., Oostra, B., LeMasson, G., et al. (2014). Dendritic channelopathies contribute to neocortical and sensory hyperexcitability in Fmr1(-/y) mice. Nat Neurosci 17, 1701–1709. 10.1038/nn.3864.

29. Chen, Q., Deister, C.A., Gao, X., Guo, B., Lynn-Jones, T., Chen, N., Wells, M.F., Liu, R., Goard, M.J., Dimidschstein, J., et al. (2020). Dysfunction of cortical GABAergic neurons leads to sensory hyper-reactivity in a Shank3 mouse model of ASD. Nat Neurosci 23, 520–532. 10.1038/s41593-020-0598-6.

30. Goel, A., Cantu, D.A., Guilfoyle, J., Chaudhari, G.R., Newadkar, A., Todisco, B., de Alba, D., Kourdougli, N., Schmitt, L.M., Pedapati, E., et al. (2018). Impaired perceptual learning in a mouse model of Fragile X syndrome is mediated by parvalbumin neuron dysfunction and is reversible. Nat Neurosci 21, 1404– 1411. 10.1038/s41593-018-0231-0.

31. Wallace, M.L., van Woerden, G.M., Elgersma, Y., Smith, S.L., and Philpot, B.D. (2017). Ube3a loss increases excitability and blunts orientation tuning in the visual cortex of Angelman syndrome model mice. J Neurophysiol 118, 634–646. 10.1152/jn.00618.2016.

32. Juczewski, K., von Richthofen, H., Bagni, C., Celikel, T., Fisone, G., and Krieger, P. (2016). Somatosensory map expansion and altered processing of tactile inputs in a mouse model of fragile X syndrome. Neurobiol Dis SC, 201–215. 10.1016/j.nbd.2016.09.007.

33. Rotschafer, S., and Razak, K. (2013). Altered auditory processing in a mouse model of fragile X syndrome. Brain Res 1506, 12–24. 10.1016/j.brainres.2013.02.038.

34. He, C.X., Cantu, D.A., Mantri, S.S., Zeiger, W.A., Goel, A., and Portera-Cailliau, C. (2017). Tactile Defensiveness and Impaired Adaptation of Neuronal Activity in the Fmr1 Knock-Out Mouse Model of Autism. J Neurosci 37, 6475–6487. 10.1523/JNEUROSCI.0651-17.2017.

35. Gandal, M.J., Edgar, J.C., Ehrlichman, R.S., Mehta, M., Roberts, T.P.L., and Siegel, S.J. (2010). Validating γ oscillations and delayed auditory responses as translational biomarkers of autism. Biol Psychiatry 68, 1100–1106. 10.1016/j.biopsych.2010.09.031.

36. Lovelace, J.W., Ethell, I.M., Binder, D.K., and Razak, K.A. (2018). Translation-relevant EEG phenotypes in a mouse model of Fragile X Syndrome. Neurobiol Dis 115, 39–48. 10.1016/j.nbd.2018.03.012.

37. Monday, H.R., Wang, H.C., and Feldman, D.E. (2023). Circuit-level theories for sensory dysfunction in autism: convergence across mouse models. Front. Neurol. 14. 10.3389/fneur.2023.1254297.

38. Simons, D.J., and Woolsey, T.A. (1979). Functional organization in mouse barrel cortex. Brain Research 165, 327–332. 10.1016/0006-8993(79)90564-X.

39. Petersen, C.C.H. (2019). Sensorimotor processing in the rodent barrel cortex. Nat Rev Neurosci 20, 533–546. 10.1038/s41583-019-0200-y.

40. Woolsey, T.A., and Van der Loos, H. (1970). The structural organization of layer IV in the somatosensory region (SI) of mouse cerebral cortex. The description of a cortical field composed of discrete cytoarchitectonic units. Brain Res 17, 205–242. 10.1016/0006-8993(70)90079-x.

41. Orefice, L.L., Zimmerman, A.L., Chirila, A.M., Sleboda, S.J., Head, J.P., and Ginty, D.D. (2016). Peripheral Mechanosensory Neuron Dysfunction Underlies Tactile and Behavioral Deficits in Mouse Models of ASDs. Cell 166, 299–313. 10.1016/j.cell.2016.05.033.

42. Gonçalves, J.T., Anstey, J.E., Golshani, P., and Portera-Cailliau, C. (2013). Circuit level defects in the developing neocortex of Fragile X mice. Nat Neurosci 16, 903–909. 10.1038/nn.3415.

43. Kourdougli, N., Suresh, A., Liu, B., Juarez, P., Lin, A., Chung, D.T., Graven Sams, A., Gandal, M.J., Martínez-Cerdeño, V., Buonomano, D.V., et al. (2023). Improvement of sensory deficits in fragile X mice by increasing cortical interneuron activity after the critical period. Neuron 111, 2863–2880.e6. 10.1016/j.neuron.2023.06.009.

44. Li, M., Eltabbal, M., Tran, H.-D., and Kuhn, B. (2023). Scn2a insufficiency alters spontaneous neuronal Ca2+ activity in somatosensory cortex during wakefulness. iScience 26, 108138. 10.1016/j.isci.2023.108138.

45. Wang, H.C., and Feldman, D.E. (2024). Degraded tactile coding in the Cntnap2 mouse model of autism. Cell Rep 43, 114612. 10.1016/j.celrep.2024.114612.

46. Matharu, N., Rattanasopha, S., Tamura, S., Maliskova, L., Wang, Y., Bernard, A., Hardin, A., Eckalbar, W.L., Vaisse, C., and Ahituv, N. (2019). CRISPR-mediated activation of a promoter or enhancer rescues obesity caused by haploinsufficiency. Science 363, eaau0629. 10.1126/science.aau0629.

47. Tamura, S., Nelson, A.D., Spratt, P.W.E., Hamada, E.C., Zhou, X., Kyoung, H., Li, Z., Arnould, C., Barskyi, V., Krupkin, B., et al. (2025). CRISPR activation for SCN2A-related neurodevelopmental disorders. Nature. 10.1038/s41586-025-09522-w.

48. Antoine, M.W., Langberg, T., Schnepel, P., and Feldman, D.E. (2019). Increased Excitation-Inhibition Ratio Stabilizes Synapse and Circuit Excitability in Four Autism Mouse Models. Neuron 101, 648–661.e4. 10.1016/j.neuron.2018.12.026.

49. Monday, H.R., Wang, H.C., and Feldman, D.E. (2023). Circuit-level theories for sensory dysfunction in autism: convergence across mouse models. Front Neurol 14, 1254297. 10.3389/fneur.2023.1254297.

50. Wang, H.C., and Feldman, D.E. (2023). Degraded Tactile Coding in the Cntnap2 Mouse Model of Autism. Preprint, 10.2139/ssrn.4620291 https://doi.org/10.2139/ssrn.4620291.

51. Goel, A., Cantu, D.A., Guilfoyle, J., Chaudhari, G.R., Newadkar, A., Todisco, B., de Alba, D., Kourdougli, N., Schmitt, L.M., Pedapati, E., et al. (2018). Impaired perceptual learning in a mouse model of Fragile X syndrome is mediated by parvalbumin neuron dysfunction and is reversible. Nat Neurosci 21, 1404– 1411. 10.1038/s41593-018-0231-0.

52. Wadle, S.L., Ritter, T.C., Wadle, T.T.X., and Hirtz, J.J. (2024). Topography and Ensemble Activity in the Auditory Cortex of a Mouse Model of Fragile X Syndrome. eNeuro 11. 10.1523/ENEURO.0396-23.2024.

53. Brecht, M., and Sakmann, B. (2002). -Dynamic representation of whisker deflection by synaptic potentials in spiny stellate and pyramidal cells in the barrels and septa of layer 4 rat somatosensory cortex. The Journal of Physiology 543, 49–70. 10.1113/jphysiol.2002.018465.

54. Moore, C.I., and Nelson, S.B. (1998). Spatio-Temporal Subthreshold Receptive Fields in the Vibrissa Representation of Rat Primary Somatosensory Cortex. Journal of Neurophysiology 80, 2882–2892. 10.1152/jn.1998.80.6.2882.

55. De Kock, C.P.J., Bruno, R.M., Spors, H., and Sakmann, B. (2007). Layer- and cell-type-specific suprathreshold stimulus representation in rat primary somatosensory cortex. The Journal of Physiology 581, 139–154. 10.1113/jphysiol.2006.124321.

56. Vinje, W.E., and Gallant, J.L. (2000). Sparse coding and decorrelation in primary visual cortex during natural vision. Science 287, 1273–1276. 10.1126/science.287.5456.1273.

57. Drew, P.J., and Feldman, D.E. (2009). Intrinsic Signal Imaging of Deprivation-Induced Contraction of Whisker Representations in Rat Somatosensory Cortex. Cereb Cortex 16, 331–348. 10.1093/cercor/bhn085.

58. Frostig, R.D., and Chen-Bee, C.H. (2009). Visualizing Adult Cortical Plasticity Using Intrinsic Signal Optical Imaging. In In Vivo Optical Imaging of Brain Function Frontiers in Neuroscience., R. D. Frostig, ed. (CRC Press/Taylor C Francis).

59. Grinvald, A., Lieke, E., Frostig, R.D., Gilbert, C.D., and Wiesel, T.N. (1986). Functional architecture of cortex revealed by optical imaging of intrinsic signals. Nature 324, 361–364. 10.1038/324361a0.

60. Masino, S.A., and Frostig, R.D. (1996). Quantitative long-term imaging of the functional representation of a whisker in rat barrel cortex. Proceedings of the National Academy of Sciences S3, 4942–4947. 10.1073/pnas.93.10.4942.

61. Buzsáki, G., Anastassiou, C.A., and Koch, C. (2012). The origin of extracellular fields and currents — EEG, ECoG, LFP and spikes. Nat Rev Neurosci 13, 407–420. 10.1038/nrn3241.

62. Destexhe, A., Contreras, D., and Steriade, M. (1999). Spatiotemporal analysis of local field potentials and unit discharges in cat cerebral cortex during natural wake and sleep states. J Neurosci 19, 4595–4608. 10.1523/JNEUROSCI.19-11-04595.1999.

63. Lee, J.-H., Shin, H.-S., Lee, K.-H., and Chung, S. (2015). LFP-guided targeting of a cortical barrel column for in vivo two-photon calcium imaging. Sci Rep 5, 15905. 10.1038/srep15905.

64. Trees, H.L.V. (2004). Detection, Estimation, and Modulation Theory, Part I: Detection, Estimation, and Linear Modulation Theory (John Wiley C Sons).

65. Nelson, A.D., Catalfio, A.M., Gupta, J.P., Min, L., Caballero-Florán, R.N., Dean, K.P., Elvira, C.C., Derderian, K.D., Kyoung, H., Sahagun, A., et al. (2024). Physical and functional convergence of the autism risk genes Scn2a and Ank2 in neocortical pyramidal cell dendrites. Neuron 112, 1133–1149.e6. 10.1016/j.neuron.2024.01.003.

66. Stern, E.A., Maravall, M., and Svoboda, K. (2001). Rapid development and plasticity of layer 2/3 maps in rat barrel cortex in vivo. Neuron 31, 305–315. 10.1016/s0896-6273(01)00360-9.

67. Lendvai, B., Stern, E.A., Chen, B., and Svoboda, K. (2000). Experience-dependent plasticity of dendritic spines in the developing rat barrel cortex in vivo. Nature 404, 876–881. 10.1038/35009107.

68. Maravall, M., Stern, E.A., and Svoboda, K. (2004). Development of intrinsic properties and excitability of layer 2/3 pyramidal neurons during a critical period for sensory maps in rat barrel cortex. J Neurophysiol 31, 144–156. 10.1152/jn.00598.2003.

69. Wen, J.A., and Barth, A.L. (2011). Input-specific critical periods for experience-dependent plasticity in layer 2/3 pyramidal neurons. J Neurosci 31, 4456–4465. 10.1523/JNEUROSCI.6042-10.2011.

70. Wang, C., Derderian, K.D., Hamada, E., Zhou, X., Nelson, A.D., Kyoung, H., Ahituv, N., Bouvier, G., and Bender, K.J. (2024). Impaired cerebellar plasticity hypersensitizes sensory reflexes in SCN2A-associated ASD. Neuron 112, 1444–1455.e5. 10.1016/j.neuron.2024.01.029.

71. Tamura, S., Nelson, A.D., Spratt, P.W.E., Kyoung, H., Zhou, X., Li, Z., Zhao, J., Holden, S.S., Sahagun, A., Keeshen, C.M., et al. (2022). CRISPR activation rescues abnormalities in SCN2A haploinsufficiency-associated autism spectrum disorder. Preprint at bioRxiv, 10.1101/2022.03.30.486483 https://doi.org/10.1101/2022.03.30.486483.

72. Matharu, N., Rattanasopha, S., Tamura, S., Maliskova, L., Wang, Y., Bernard, A., Hardin, A., Eckalbar, W.L., Vaisse, C., and Ahituv, N. (2019). CRISPR-mediated activation of a promoter or enhancer rescues obesity caused by haploinsufficiency. Science 363, eaau0629. 10.1126/science.aau0629.

73. Lakens, D. (2017). Equivalence Tests: A Practical Primer for t Tests, Correlations, and Meta-Analyses. Soc Psychol Personal Sci 8, 355–362. 10.1177/1948550617697177.

74. Kourdougli, N., Suresh, A., Liu, B., Juarez, P., Lin, A., Chung, D.T., Graven Sams, A., Gandal, M.J., Martínez-Cerdeño, V., Buonomano, D.V., et al. (2023). Improvement of sensory deficits in fragile X mice by increasing cortical interneuron activity after the critical period. Neuron 111, 2863–2880.e6. 10.1016/j.neuron.2023.06.009.

75. Deemyad, T., Puig, S., Papale, A.E., Qi, H., LaRocca, G.M., Aravind, D., LaNoce, E., and Urban, N.N. (2022). Lateralized Decrease of Parvalbumin+ Cells in the Somatosensory Cortex of ASD Models Is Correlated with Unilateral Tactile Hypersensitivity. Cereb Cortex 32, 554–568. 10.1093/cercor/bhab233.

76. Vormstein-Schneider, D., Lin, J.D., Pelkey, K.A., Chittajallu, R., Guo, B., Arias-Garcia, M.A., Allaway, K., Sakopoulos, S., Schneider, G., Stevenson, O., et al. (2020). Viral manipulation of functionally distinct interneurons in mice, non-human primates and humans. Nat Neurosci 23, 1629–1636. 10.1038/s41593-020-0692-9.

77. Gainey, M.A., and Feldman, D.E. (2017). Multiple shared mechanisms for homeostatic plasticity in rodent somatosensory and visual cortex. Philos Trans R Soc Lond B Biol Sci 372, 20160157. 10.1098/rstb.2016.0157.

78. Gainey, M.A., Aman, J.W., and Feldman, D.E. (2018). Rapid Disinhibition by Adjustment of PV Intrinsic Excitability during Whisker Map Plasticity in Mouse S1. J. Neurosci. 38, 4749–4761. 10.1523/JNEUROSCI.3628-17.2018.

79. Li, L., Gainey, M.A., Goldbeck, J.E., and Feldman, D.E. (2014). Rapid homeostasis by disinhibition during whisker map plasticity. Proceedings of the National Academy of Sciences of the United States of America 111, 1616–1621.

80. Monday, H.R., Nieto, A.M., Yohannes, S.A., Luxu, S., Wong, K.W., Bolio, F.E., and Feldman, D.E. (2025). Physiological and molecular impairment of PV circuit homeostasis in mouse models of autism. Preprint, 10.1101/2025.01.08.632056 https://doi.org/10.1101/2025.01.08.632056.

81. Wallace, M.L., van Woerden, G.M., Elgersma, Y., Smith, S.L., and Philpot, B.D. (2017). Ube3a loss increases excitability and blunts orientation tuning in the visual cortex of Angelman syndrome model mice. J Neurophysiol 118, 634–646. 10.1152/jn.00618.2016.

82. Banerjee, A., Rikhye, R.V., Breton-Provencher, V., Tang, X., Li, C., Li, K., Runyan, C.A., Fu, Z., Jaenisch, R., and Sur, M. (2016). Jointly reduced inhibition and excitation underlies circuit-wide changes in cortical processing in Rett syndrome. Proc Natl Acad Sci U S A 113, E7287–E7296. 10.1073/pnas.1615330113.

83. Michaelson, S.D., Ozkan, E.D., Aceti, M., Maity, S., Llamosas, N., Weldon, M., Mizrachi, E., Vaissiere, T., Gaffield, M.A., Christie, J.M., et al. (2018). SYNGAP1 Heterozygosity Disrupts Sensory Processing by Reducing Touch-Related Activity within Somatosensory Cortex Circuits. Nat Neurosci 21, 1–13. 10.1038/s41593-018-0268-0.

84. Gauld, O.M., Packer, A.M., Russell, L.E., Dalgleish, H.W.P., Iuga, M., Sacadura, F., Roth, A., Clark, B.A., and Häusser, M. (2024). A latent pool of neurons silenced by sensory-evoked inhibition can be recruited to enhance perception. Neuron 112, 2386–2403.e6. 10.1016/j.neuron.2024.04.015.

85. O’Connor, D.H., Hires, S.A., Guo, Z.V., Li, N., Yu, J., Sun, Q.-Q., Huber, D., and Svoboda, K. (2013). Neural coding during active somatosensation revealed using illusory touch. Nature neuroscience 16, 958–965.

86. Zhang, J., Chen, X., Eaton, M., Wu, J., Ma, Z., Lai, S., Park, A., Ahmad, T.S., Que, Z., Lee, J.H., et al. (2021). Severe deficiency of the voltage-gated sodium channel NaV1.2 elevates neuronal excitability in adult mice. Cell Reports 36. 10.1016/j.celrep.2021.109495.

87. Turrigiano, G.G., and Nelson, S.B. (2004). Homeostatic plasticity in the developing nervous system. Nature reviews. Neuroscience 5, 97–107.

88. Hensch, T.K. (2005). Critical period plasticity in local cortical circuits. Nature reviews. Neuroscience 6, 877–888.

89. Lee, S.-H., Kwan, A.C., Zhang, S., Phoumthipphavong, V., Flannery, J.G., Masmanidis, S.C., Taniguchi, H., Huang, Z.J., Zhang, F., Boyden, E.S., et al. (2012). Activation of specific interneurons improves V1 feature selectivity and visual perception. Nature 488, 379–383. 10.1038/nature11312.

90. Yoo, Y.-E., Mandal, P., Tang, Z., Zhang, Z., Zhang, J., Chen, X., Robinson, M., Eaton, M., Deming, B., Halurkar, M., et al. (2025). Single-nucleus transcriptomics reveal disrupted pathways in the prefrontal cortex of Scn2a -deficient mice. Preprint, 10.1101/2025.09.16.676420 https://doi.org/10.1101/2025.09.16.676420.

91. Erzurumlu, R.S., and Gaspar, P. (2012). Development and critical period plasticity of the barrel cortex. Eur J Neurosci 35, 1540–1553. 10.1111/j.1460-9568.2012.08075.x.

92. Planells-Cases, R., Caprini, M., Zhang, J., Rockenstein, E.M., Rivera, R.R., Murre, C., Masliah, E., and Montal, M. (2000). Neuronal death and perinatal lethality in voltage-gated sodium channel alpha(II)-deficient mice. Biophys J 78, 2878–2891. 10.1016/S0006-3495(00)76829-9.

93. Finnerty, G.T., Roberts, L.S.E., and Connors, B.W. (1999). Sensory experience modifies the short-term dynamics of neocortical synapses. Nature 400, 367–371. 10.1038/22553.

94. Allen, C.B., Celikel, T., and Feldman, D.E. (2003). Long-term depression induced by sensory deprivation during cortical map plasticity in vivo. Nature neuroscience C, 291–299.

95. Pernía-Andrade, A.J., Goswami, S.P., Stickler, Y., Fröbe, U., Schlögl, A., and Jonas, P. (2012). A deconvolution-based method with high sensitivity and temporal resolution for detection of spontaneous synaptic currents in vitro and in vivo. Biophys J 103, 1429–1439. 10.1016/j.bpj.2012.08.039.

96. Pachitariu, M., Steinmetz, N., Kadir, S., Carandini, M., and D, H.K. (2016). Kilosort: realtime spike-sorting for extracellular electrophysiology with hundreds of channels. bioRxiv. 10.1101/061481.

97. Mayhew, J.E.W., and Zheng, Y. (1996). A Model of the Intrinsic Image Signal and an Evaluation of the Methodology of Intrinsic Image Signal Analysis. https://eprints.whiterose.ac.uk/80348/.

98. Rossant, C., Kadir, S.N., Goodman, D.F.M., Schulman, J., Hunter, M.L.D., Saleem, A.B., Grosmark, A., Belluscio, M., Denfield, G.H., Ecker, A.S., et al. (2016). Spike sorting for large, dense electrode arrays. Nat. Neurosci. 19, 634–641.

